# No-Go Decay mRNA cleavage in the ribosome exit tunnel produces 5’-OH ends phosphorylated by Trl1

**DOI:** 10.1101/465633

**Authors:** Albertas Navickas, Sébastien Chamois, Rénette Saint-Fort, Julien Henri, Claire Torchet, Lionel Benard

## Abstract

The No-Go Decay (NGD) mRNA surveillance pathway degrades mRNAs containing stacks of stalled ribosomes. Although an endoribonuclease has been proposed to initiate cleavages upstream of the stall sequence, the production of two RNA fragments resulting from a unique cleavage has never been demonstrated. We have used mRNAs expressing a 3’-ribozyme to produce truncated transcripts *in vivo* to mimic naturally occurring truncated mRNAs known to trigger NGD. This technique allows us to analyse endonucleolytic cleavage events at single-nucleotide resolution starting at the third collided ribosome, which we show to be Hel2-dependent. These cleavages map precisely in the mRNA exit tunnel of the ribosome, 8 nucleotides upstream of the first P-site residue and release 5’-hydroxylated RNA fragments requiring 5’-phosphorylation prior to digestion by the exoribonuclease Xrn1, or alternatively by Dxo1. Finally, we identify the RNA kinase Trl1, alias Rlg1, as an essential player in the degradation of NGD RNAs.

## Introduction

The No-Go Decay (NGD) mRNA surveillance pathway degrades mRNAs containing stalled ribosomes^1, 2^. NGD occurs when translation elongation is blocked by the presence of stable intra- or intermolecular RNA structures, enzymatic cleavage, chemically damaged sequences or rare codons^1, 3–8^. This mRNA degradation process is dependent on translation and involves an endoribonuclease that cleaves just upstream of the stall sequence^1, 5, 6, 9^. Other mRNA surveillance pathways can also ultimately lead to NGD. For instance, transcripts synthesized without a stop codon due to premature polyadenylation have stalled ribosomes that are initially detected by the Non-Stop Decay (NSD) pathway^9, 10^. NSD targeted mRNAs are cleaved by an uncharacterized mechanism and become targets of NGD when ribosomes reach the new 3’-end and stall^9, 11, 12^. NGD thus plays a key role in resolving translational issues potentially detrimental to cellular homeostasis. When mRNAs are truncated, the stalled ribosomes are rescued in a process mediated by the Dom34/Hbs1 complex that dissociates the ribosomal subunits^5^. Their association with the 60S subunit is recognized by the Ribosome Quality Control (RQC) pathway, leading to the rapid degradation of the nascent peptide^13–15^. However, despite extensive study, the precise location of NGD cleavage and the mechanism of degradation of the resulting RNA fragment remain elusive.

In this paper, we focus on the fate of NGD-cleaved mRNAs, with an initial goal of mapping the sites of mRNA cleavage with accuracy. Two major obstacles to achieving this objective are that NGD fragments are rapidly attacked by 5’-3’ and 3’-5’ exoribonucleases after ribosome dissociation^5^ and that simultaneously blocking the 5’-3’ and 3’-5’ exoribonuclease decay pathways is synthetically lethal^16^. It has been shown, however, that the stability of such mRNAs is largely dependent on the Dom34/Hbs1 complex^5, 17^. In *dom34* mutant cells, ribosomes stalled at the 3’-end of truncated mRNAs inhibit the degradation by the exosome and facilitate the detection of sequential endonucleolytic cleavages upstream of the ribosomal stall site^5^. Interestingly, *dom34* and *xrn1* mutations (inactivating the main 5’-3’ exonucleolytic degradation pathway) are not synthetic lethal^1^. Moreover, NGD endonucleolytic cleavages still occur in the absence of Dom34^2, 3^. The limited 3’-5’ degradation of specific mRNA targets (in the absence of Dom34) combined with 5’-3’ exoribonuclease mutants thus allows an accumulation of RNA fragments resulting from endonucleolytic cleavages whose extremities can be mapped accurately. We have created truncated mRNAs *in vivo* by insertion of a hammerhead ribozyme sequence (Rz)^18^ known to generate NGD targeted mRNAs^5^. This construction mimics chemically or enzymatically cleaved mRNAs, or those resulting from abortively spliced mRNAs that are processed by the NGD pathway^5, 9, 19^. As anticipated, these designed truncated 3’-ends block ribosomes at determined positions and, because ribosomes guide NGD mRNA cleavages^5, 20^, we were able to detect 3’-NGD RNA fragments of specific sizes. By analysing these RNAs in detail, we show the importance of the 5’-3’ exoribonuclease Xrn1 in the production of 3’-NGD fragments. We also perfectly match a 3’-NGD cleavage product with a 5’-NGD cleavage fragment in the region of the third stalled ribosome. We have mapped this site and show that a unique endonucleolytic cleavage occurs 8 nucleotides (nts) upstream of the first P-site nt within the third stacked ribosome. The two leading ribosomes are apparently not competent for this cleavage. We demonstrate that this 3’-NGD RNA has a hydroxylated 5’-extremity and show that 5’-phosphorylation by the Trl1 kinase^21^ is required to allow degradation by 5’-3’ exoribonucleases. Interestingly, in the absence of Xrn1, the alternative 5’-3’ exoribonuclease Dxo1 takes over^22^. We additionally analysed mRNAs containing rare codons and demonstrate that at least three stacked ribosomes are also required for endonucleolytic cleavage of these mRNAs. We show that 5’ ends observed in regions covered by disomes are the result of 5’-3’ trimming by Dxo1.

## Results

### Mapping the 5’-ends of 3’-NGD RNA fragments

To generate 3’-truncated mRNA substrates for NGD *in vivo*, we inserted a hammerhead ribozyme sequence^18^ in the 3’-sequence of the *URA3* gene ORF (mRNA1Rz). This results in the production of an mRNA that lacks a stop codon and a polyadenylated tail, called mRNA1 in Fig. 1a and Supplementary Fig. 1a, and that is known to be an NGD target^5^. We first verified that we could detect NGD cleavages in the 3’-proximal region of mRNA1, by northern blotting with a probe corresponding to the 3’-end (probe prA, Fig. 1a and Supplementary Fig. 1a). The upstream and downstream cleavage products are referred to as 5’-NGD and 3’-NGD RNAs, respectively (Fig. 1a). We indeed detected a ladder of 3’-NGD RNA fragments in *dom34* mutant cells (Fig. 1b), in the presence or absence of active 5’-3’ or 3’-5’ exonucleolytic decay pathways, *i.e. xrn1* or *ski2* mutations, respectively^23^. In agreement with the current NGD model in which endonucleolytically cleaved 3’-NGD fragments are primarily degraded by the 5’-3’ exoribonuclease Xrn1^5^, inactivation of the 5’-3’ RNA decay pathway (*xrn1* mutant cells) produced a different ladder of 3’-NGD RNAs compared to WT or the *ski2* mutant. This was confirmed by a higher resolution PAGE analysis followed by northern blotting (Fig. 1c). The PAGE analysis was completed by mapping the 5’-ends of the 3’-NGD RNA fragments in the *dom34* and *dom34/xrn1* mutants by primer extension experiments with prA (Fig. 1d). We showed that the truncated mRNAs produce several discrete 3’-NGD RNA bands (B1 to B5) that can be mapped to single-nucleotide resolution. B5 (77 nts) and the major RNA species B1 (47 nts) visible in the *dom34* mutant were not detected in the absence of the 5’-3’ exoribonuclease Xrn1 (Fig. 1b-d). B3 (68 nts) and B2 (65 nts) RNAs were exclusively observed in the *xrn1* mutant cells, and B4 (71 nts) was detected in all four strains (Fig. 1c). In addition, we analysed a *dcp2Δ* decapping mutant that totally blocks 5’-3’ exoribonucleolytic attacks on the 5’-end of mRNAs (Fig. 1b, c)^23, 24^. The particularly strong growth defect of the *dcp2Δ* mutant may explain the decrease in the amount of NGD RNA detected but, interestingly, the fact that NGD RNAs remained detectable (Fig. 1c) suggests that an alternative pathway bypasses mRNA decapping for their production.

**Fig. 1.**
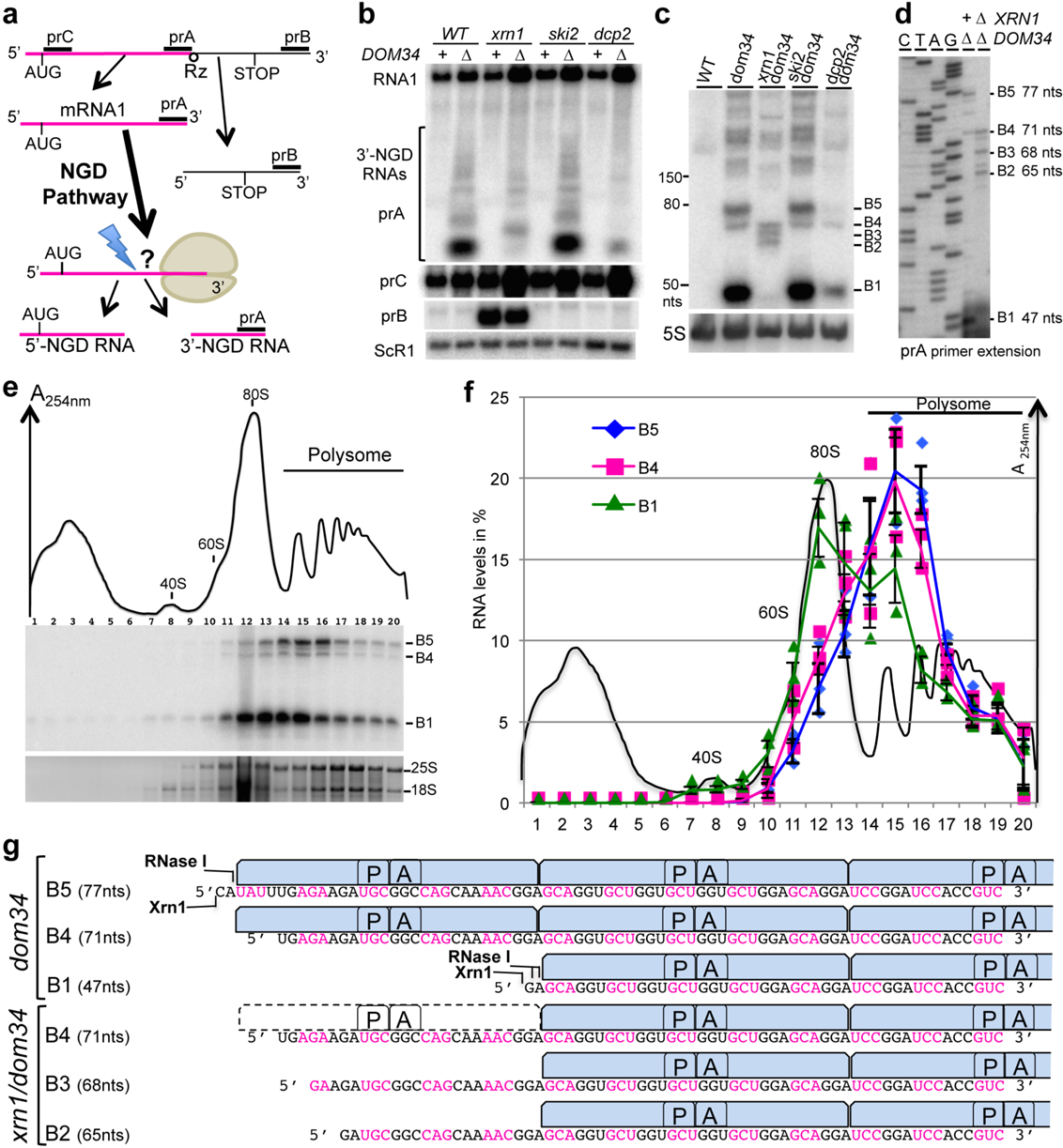
Size characterization of 3’-NGD RNA fragments and ribosomal association. **a** Schematic view of the URA3Rz mRNA showing the ribozyme (Rz) site (see also Supplementary Fig. 1a). Translational start (AUG) and stop codons are indicated. RNA1 (in magenta) is the stop-codon-less mRNA following ribozyme cleavage. Probes prA, prB and prC used in northern blots analysis are indicated. 5’ and 3’-NGD RNAs are the products of NGD cleavage of mRNA1. The lightning flash represents the NGD endonucleolytic cleavage and probe prA is designed for the detection of all potential 3’-NGD RNAs (see also Supplementary Fig. 1a). **b** Agarose gel electrophoresis followed by northern blot showing levels of mRNA1 and 3’-NGD RNA fragments in the indicated strains. The ScR1 RNA served as a loading control**. c** Analysis similar to (**b**) using 8% PAGE. The 5S rRNA served as a loading control. **d** Primer extension experiments using probe prA to determine the 5’-end of 3’-NGD RNAs. B1, B2, B3, B4 and B5 RNAs shown in Fig. 1c are indicated with the corresponding size in nucleotides (nts) calculated by primer extension. **e** Analysis of 3’-NGD RNA association with ribosomes in *dom34* mutant cells. Upper, on ribosome profile, positions of 40S and 60S subunits, 80S and polysome are indicated. Panel below, 20 fractions were collected and extracted RNAs were analysed similarly to (**c**). Bottom panel: 18S and 25S rRNAs are shown as references for ribosome and polysome sedimentation, using 1% agarose gel electrophoresis and ethidium bromide staining. **f** Distribution of the 3’-NGD RNAs analysed in (**e**). For each fraction, levels of B1, B4 and B5 RNAs were plotted as a % of total amount of B1, B4 and B5 RNAs respectively. The profile in (**e**) is reported in (**f**). **g** Schematic view of the ribosome positioning on 3’-NGD RNAs combining information on size resolution, ribosomal association (Fig. 1), and RNase ribosomal protection of 3’-NGD RNAs (Supplementary Fig. 1). Codons are shown in black or magenta. A and P are ribosome A-and P-sites respectively. Error bars indicate standard deviation (s.d) calculated from three independent experiments. Source data are provided as a Source Data file.

The sizes of the major B1 and B5 RNAs differ by 30 nts (Fig. 1d), consistent with the length of mRNA covered by an individual ribosome^25^. We therefore surmised that the difference in size is most likely due to the presence of an extra ribosome protecting the B5 RNA species from 5’-3’ degradation by Xrn1, compared to B1^26^. This prompted us to analyse the association of these 3’-NGD RNAs with ribosomes in sucrose gradients.

### Ribosome association with 3’-NGD RNA fragments

We performed polysome analyses to assess the distribution of 3’-NGD RNAs in different ribosomal fractions in *dom34* mutant cells (Fig. 1e, f). The B1 RNA (47 nts) was found to associate with monosomes (fractions 11 to 13) and disomes (fraction 15), with disomes being the theorical maximal coverage of such a short RNA, as proposed by^9, 20, 27^. Indeed, the association of a 47-nt RNA species with disomes has been deduced from ribosome profiling experiments^20^ and is explained by the approximate size of the trailing ribosome protecting a full ribosome footprint (28–30 nt) and the leading ribosome protecting a half footprint to the site of mRNA truncation (16–17 nts, with no RNA or an incomplete codon in the A-site). An additional ribosome would thus be expected to protect a ∼77-nt RNA. Accordingly, the B5 RNA (77 nts) was found to associate with two and three ribosomes, in fractions 15 and 16, respectively. The 71-nt B4 RNA was also associated with the disome and trisome peaks (Fig. 1e, f). We suspect that ribosomes do not stay stably bound to the different RNA species during sucrose gradient analysis, but this experiment allows us to propose that two ribosomes maximally associate with B1, while up to three can associate with B4 and B5 *in vivo*. This also prompted us to determine how these 3’-NGD RNAs were protected from Xrn1 and RNase I activities in cell extracts prior to centrifugation on sucrose cushions (Supplementary Fig. 1). These complementary experiments show that ribosomes persist and protect these RNA species from RNases *in vitro* and support results presented above (Supplementary Notes 1 and 2 and Supplementary Fig. 1b-i).

The results in Fig. 1 and Supplementary Fig. 1 allow us to infer the precise positions of ribosomes on B5 and B4 RNA species in the *dom34* mutant backgrounds. Our experiments all converge to the conclusion that the B5 species corresponds to RNAs covered by trisomes (Fig. 1g). We also conclude that three ribosomes cover the 71-nt B4 RNA in *dom34* mutant cell extracts as this species is resistant to Xrn1 (Supplementary Note 1), its 5’-region is protected from RNase I digestion *in vitro* (Supplementary Note 2) and a significant proportion remains associated with fractions corresponding to three ribosomes in sucrose gradients (Fig. 1e, f).

### Dxo1 trimming of 3’-NGD fragments in Xrn1 deficient cells

We strongly suspected that the B4 species was the original NGD product, and because B3 and B2 RNAs were exclusively detected in Xrn1 deficient cells, we speculated that these RNAs might be derived from B4 by an alternative 5’-3’ exoribonuclease. We therefore asked whether the 5’-3’ exoribonucleolytic activity of Dxo1, which plays an important role in 5’-end capping quality control^22^, might explain the presence of the B3 and B2 RNAs. Remarkably, deletion of both *XRN1* and *DXO1* genes in a *dom34* background completely abolished the production of the B3 and B2 RNAs, and only the B4 3’-NGD species remained detectable by northern blot analysis (Fig. 2a) or in primer extension assays (Supplementary Fig. 2a). Complementation of the *dom34/xrn1/dxo1* mutant with wild-type Dxo1 restored B3 and B2 RNA production to a significant extent, but a catalytic mutant failed to do so (Fig. 2b). We took advantage of the almost exclusive presence of the B4 3’-NGD RNAs in *dom34/xrn1/dxo1* mutant cells to ask how this RNA is protected by ribosomes by adding Xrn1 to cell extracts as described above (Supplementary Fig. 1b). Some of the B4 RNA was Xrn1-resistant (Supplementary Fig. 2b) in accordance with our hypothesis that a portion of this species is protected by three ribosomes in *xrn1*/*dom34* cell extracts (Fig. 1g). Interestingly, the decrease in the amount of B4 RNA was correlated with an almost equivalent increase of a 47-nt species (Supplementary Fig. 2b, c), suggesting that disomes persist on the majority of the 3’-ends of B4 RNAs in *dom34/xrn1/dxo1* cells *in vivo*. We thus propose that two populations of B4 RNAs co-exist in Xrn1 deficient cells *in vivo*, with one population covered by three ribosomes, especially in *dom34* mutants, and the other only covered by two ribosomes, but having a 5’-protuding RNA extremity due to the absence of 5’-3’ exoribonucleases.

**Fig. 2.**
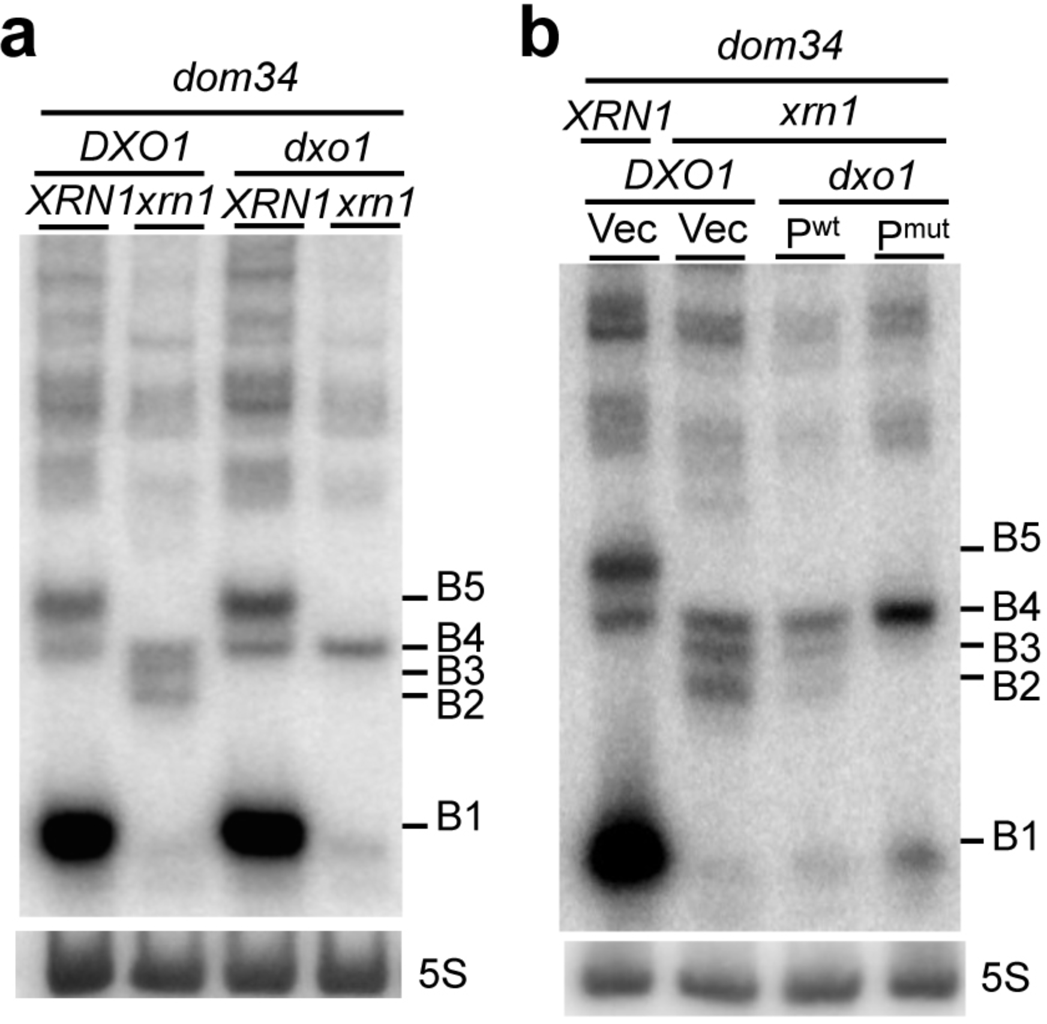
Dxo1 creates the heterogeneity of 3’-NGD RNA fragments in Xrn1 deficient cells. 8% PAGE followed by northern blot analysis using probe prA showing steady state levels of RNAs in *dom34* and other indicated mutant strains. The 5S rRNA served as a loading control**. a** Impact of *DXO1* deletion on B2 and B3 RNA production. **b** Plasmid expression of wild-type Dxo1 (P^wt^) or a Dxo1 catalytic mutant (P^mut^) (mutant E260A/D262A)^22^. The vector control is plasmid pRS313 (Vec). Source data are provided as a Source Data file.

### Mapping of the primary NGD endonucleolytic cleavage site

The results described above suggest that the principal band detected in the absence of Xrn1 and Dxo1 (B4 RNA) is a specific 3’-product of NGD cleavage in our constructs (Fig. 2a). While B4 RNA resistance to Xrn1 *in vitro* (Supplementary Fig. 1b) could be explained by a third ribosome dwelling after cleavage, we also considered the possibility that its 5’-phosphorylation state could contribute to its stability, since both Xrn1 and Dxo1 require 5’-phosphorylated extremities to degrade RNA^22, 28^. We therefore asked whether the B4 RNA naturally has a monophosphate (5’-P) or a hydroxyl group (5’-OH) at its 5’-end by treating RNA purified from *dom34* cell extracts with T4 polynucleotide kinase to see whether this would stimulate attack of B4 by Xrn1 *in vitro*. Remarkably, the B4 RNA was completely degraded by Xrn1 only after 5’-phosphorylation by T4 kinase *in vitro* (Fig. 3a), demonstrating that the B4 RNA has a 5’-OH extremity in *dom34* cells. A portion of the abundant B1 RNA persisted during Xrn1 treatment in kinase buffer (Fig. 3a, and see Methods), but parallel Xrn1 digestion in optimal buffer confirms that B1 and B5 RNAs were totally digested (*i.e*, are fully mono-phosphorylated), while the B4 RNA remained resistant (Supplementary Fig. 3a). It has recently been demonstrated that the Hel2 ubiquitin-protein ligase is crucial for the activation of NGD cleavages^29^. Consistent with our hypothesis that the B4 species corresponds to the 3’ NGD cleavage product, this RNA is no longer produced in the *hel2* mutant (Supplementary Fig. 3b).

**Fig. 3.**
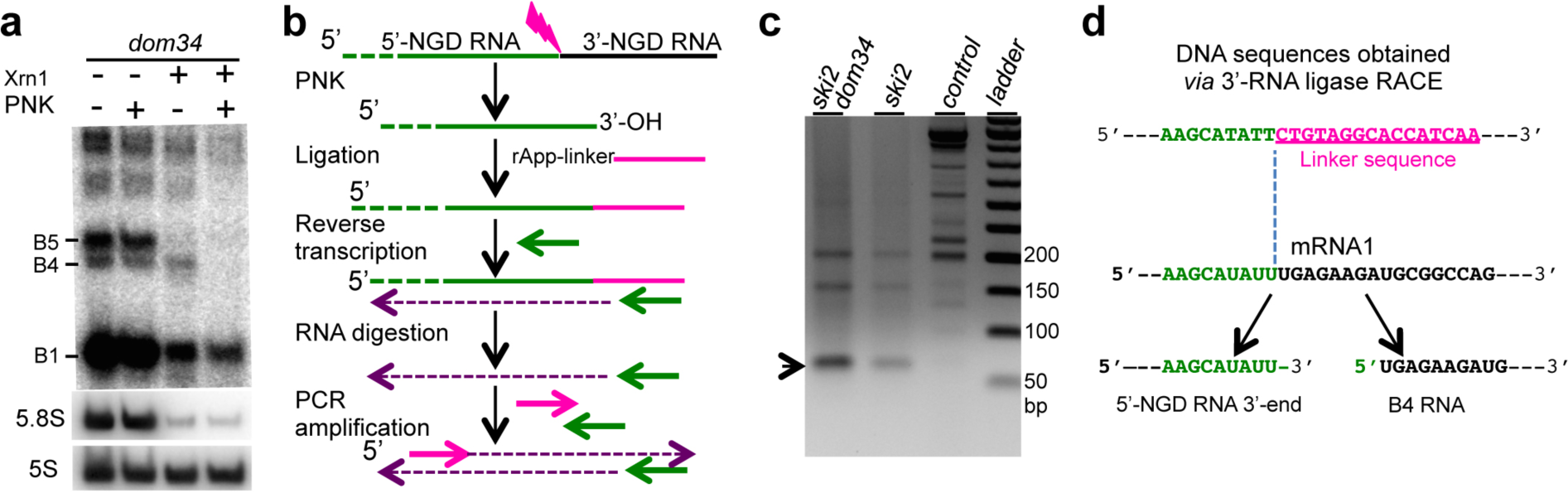
Characterization of the endonucleolytic RNA fragments. **a** Xrn1 digestion of total RNA extracts from *dom34* mutant cells in the presence or absence of polynucleotide kinase (PNK) *in vitro*. 8% PAGE followed by northern blot analysis using probe prA. The 5S rRNA served as a loading control and 5.8S rRNA as a positive control of Xrn1 treatment. **b** Flow chart illustrating the method used for 3’-RACE as described in^31^ with minor modifications according to^30^ (see Methods). **c** PCR products obtained from 3’-RACE and separated on a 2% agarose gel. Purified DNAs for sequencing are indicated by an arrowhead. Prior to PCR, cDNAs were produced from total RNA from *ski2*, *ski2/dom34* mutant cells expressing mRNA1. Control is made of total RNA from *ski2/dom34* mutant cells without mRNA1 expression. **d** Sequences obtained after 3’-RACE performed on *ski2* and *ski2/dom34* total RNA. 100% of sequenced clones (omitting a residual 5S rRNA-linker amplification detected) have this DNA sequence. 5’-NGD DNA sequence (in green) and linker sequence (in magenta). Below, the site of mRNA1 is shown before and after the cleavage producing the 3’-NGD RNA B4 and the 3’-extremity of the 5’-NGD RNA confirmed by 3’-RACE. Source data are provided as a Source Data file.

By definition endonucleolytic cleavage of RNA results in the production of 5’ and 3’-RNA fragments. However, corresponding 5’ and 3’ fragments have never been definitively demonstrated in the case of NGD targeted mRNAs. We thus searched for the corresponding 5’-NGD fragment for the B4 3’-NGD RNA. To map the 3’-end of 5’-NGD RNAs, total RNA preparations from *ski2* and *ski2*/*dom34* mutants were ligated to a pre-adenylated oligonucleotide linker using truncated RNA ligase to perform 3’-RACE (Fig. 3b)^30, 31^. The *ski2* mutant context was used to limit 3’-trimming of these RNAs *in vivo*, and RNAs were pre-treated with T4 polynucleotide kinase to modify 3’ phosphates or 2’-3’ cyclic phosphates to 3’-OH to permit RNA ligation^31^. The major RT-PCR product was of the expected size (66 bp; Fig. 3c and Supplementary Fig. 3c) and verified by sequencing the resulting clones (Fig. 3d). The identification of a matching 5’-NGD fragment for the B4 3’-NGD RNA, confirms that an endonucleolytic event occurred at this precise position. The same procedure performed on RNAs isolated from *ski2* mutants where Dom34 was still active yielded the same major PCR product, also verified by sequencing (Supplementary Fig. 3d). Thus, while the *dom34* mutation facilitates the detection of NGD fragments by increasing their stability, the cleavage event itself is Dom34-independent.

### The fate of 5’-NGD RNAs

We anticipated that following NGD cleavage of mRNA1, ribosomes that had initiated translation on the 5’-NGD fragments would advance to the new 3’-end and the RNA be subjected to Xrn1 trimming, similar to the process that generates B1 and B5 (Fig. 4a). Since the B4 3’-NGD RNAs were cut in the +1 reading frame (Fig. 1g), upstream ribosomes on these 5’-NGD RNAs would be expected to stall with one nucleotide in ribosome A-site (Fig. 4a and Supplementary Fig. 4) and as result produce new RNA fragments 47+1, 77+1 nts, protected by two or three ribosomes, respectively (see Supplementary Fig. 4). Indeed, in northern blots using probe prG, which is complementary to the new 3’-ends generated by NGD cleavage, we detected RNA fragments consistent with a 1-nt increase in size compared to those detected by prA on the same membrane (Fig. 4b). We mapped the 5’-ends of these new ribosome protected fragments by primer extension assays using prG (Fig. 4c and Supplementary Fig. 4). The detection of 48-nt (and 78-nt) cDNAs only in cells containing active Xrn1 (Fig. 4c) strongly suggests that the new NGD endonucleolytic products are covered by two and three ribosomes, respectively. The production of cDNAs of exactly the predicted sizes (48 and 78 nts) is an independent confirmation that the 3’-extremity of the 5’-NGD product corresponds precisely to the proposed NGD endonucleolytic cleavage site (Supplementary Fig. 4). Remarkably, the 3’-extremity of the 5’-NGD RNA was detected in the context of active 3’-5’ exonucleases, meaning that ribosomes run on and cover the 3’-extremity before any 3’-5’ attacks can occur. In summary, we propose that the B4 RNA is produced by endonucleolytic cleavage within the footprint of the third stalled ribosome and that at least two upstream ribosomes promptly protect the resulting 5’-NGD fragment from degradation by 3’-5’ exoribonucleases (Fig. 4d).

**Fig. 4.**
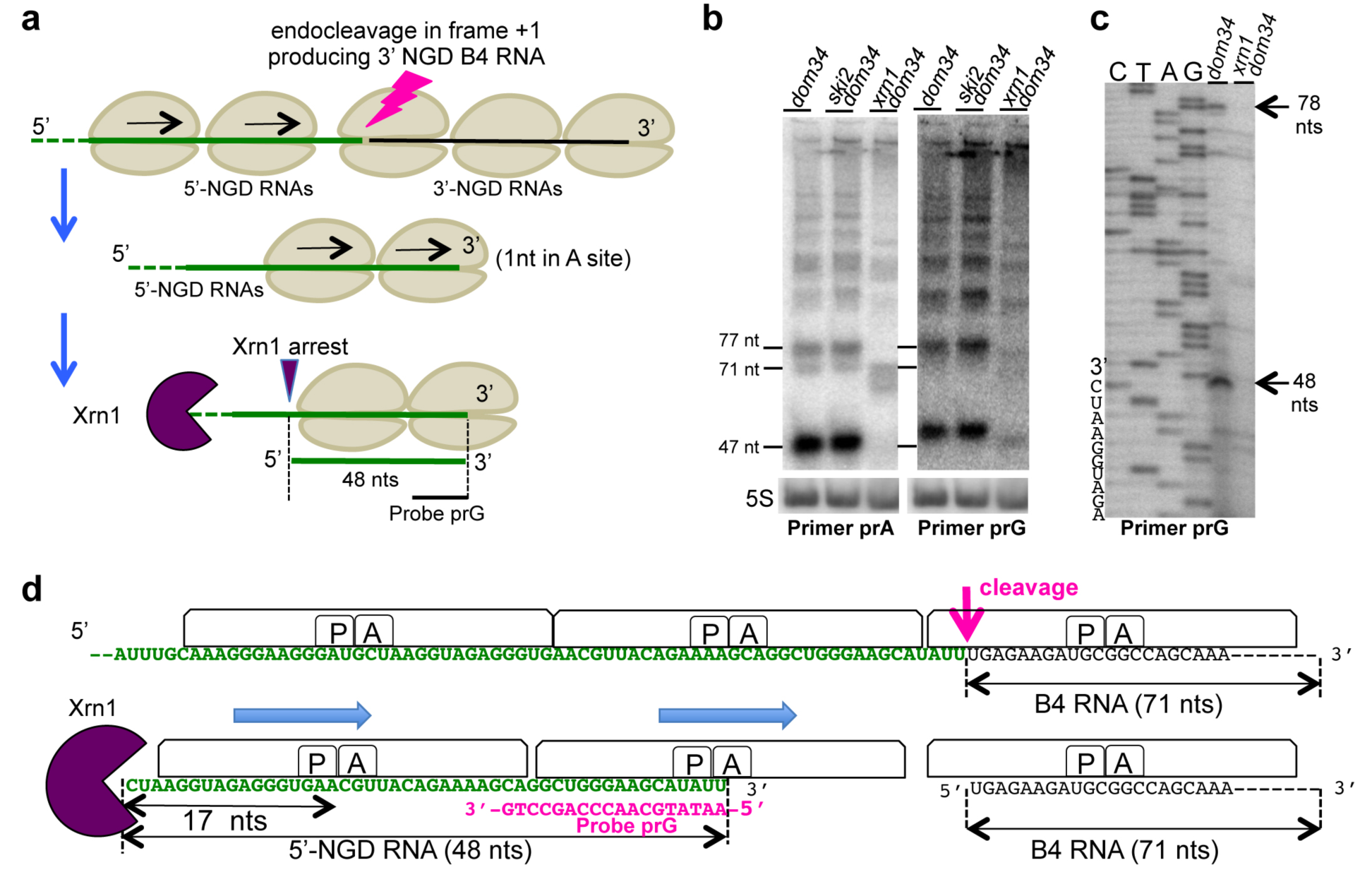
Analysis of the fate of 5’-NGD RNAs. **a** Schematic model of mRNA1 before and after the endonucleolytic cleavages producing B4 RNAs. 5’-NGD resulting RNAs are shown here covered by two ribosomes and processed by Xrn1 to 48-nt RNAs (see also Supplementary Fig. 4 for 5’-NGD RNAs covered by three ribosomes). **b** 8% PAGE followed by northern blot analysis using probe prG showing steady state levels of RNAs in the indicated mutant strains (left panel). Same membrane has been probed with prA as a ladder (right panel), and sizes of B5 (77nt), B4 (71 nt) and B1 (47nt) are indicated. The 5S rRNA served as a loading control. **c** Primer extension experiments using probe prG to determine the 5’-end of RNAs (see also Supplementary Fig. 5). **d** Schematic model of ribosome positioning on mRNA1 before and after the unique endonucleolytic cleavage producing B4 RNAs, localized 8 nts upstream of the first P-site nt. The position of disomes on the resulting 48-nt 5’-NGD RNA is shown with the distal ribosome having 1 nt in the A site (see also Supplementary Fig. 4). Source data are provided as a Source Data file.

### The 5’-OH endocleaved product is phosphorylated by Trl1

Despite the fact that a portion of the B4 species was found to be 5’-hydroxylated in *dom34* cell extracts (Fig. 3a), a major fraction of the B4 RNA from *dom34/xrn1* and *dom34*/*xrn1*/*dxo1* cell extracts can be degraded by Xrn1 *in vitro* without prior 5’– phosphorylation (Supplementary Figs 1b and 2b), suggesting that B4 accumulates as a 5’-phosphorylated species in this mutant background. A well-characterized factor with RNA kinase activity in yeast is the essential tRNA ligase Trl1^21, 32^. Splicing of tRNAs is known to generate 5’OH-intron RNAs which require Trl1 kinase activity to permit their degradation by Xrn1. We therefore asked whether the 3’-NGD B4 RNA fragments were substrates of Trl1. Since *TRL1* is an essential gene, we used *trl1Δ* strains expressing pre-spliced intron-less versions of the 10 intron-containing tRNAs that restore viability^33, 34^. If Trl1 were required for B4 degradation, loss of Trl1 function should increase the amount of B4 RNA. Remarkably, in *trl1* versus *TRL1* cells, we observed a 24-fold accumulation of the B4 RNA species and only a 2-fold-decrease of B1 RNA levels, while the B5 RNA was barely detectable (Fig. 5a, b). We asked whether the accumulated B4 RNA had a hydroxyl group (5’-OH) at its 5’-end. RNAs were analysed by 12% PAGE allowing separation of 5’-hydroxylated from 5’-phosphorylated RNAs^21^. B4 RNA migrated as two bands in *TRL1* cells whereas only the lower band was detected in *trl1Δ* cells. This species was fully shifted to the upper band after 5’-phosphorylation of RNA *in vitro* (Fig. 5c). We also verified that the B4 RNA was resistant to Xrn1 treatment *in vitro*. In contrast to B1 and B5 RNAs (Supplementary Fig. 5), B4 was completely degraded by Xrn1 only after 5’-phosphorylation by T4 kinase *in vitro* (Fig. 5d). We conclude that the B4 RNA accumulates as a fully 5’-OH species in the *trl1Δ* mutant. Thus, Trl1 is the major kinase involved in phosphorylating the B4 RNA following NGD cleavage.

**Fig. 5.**
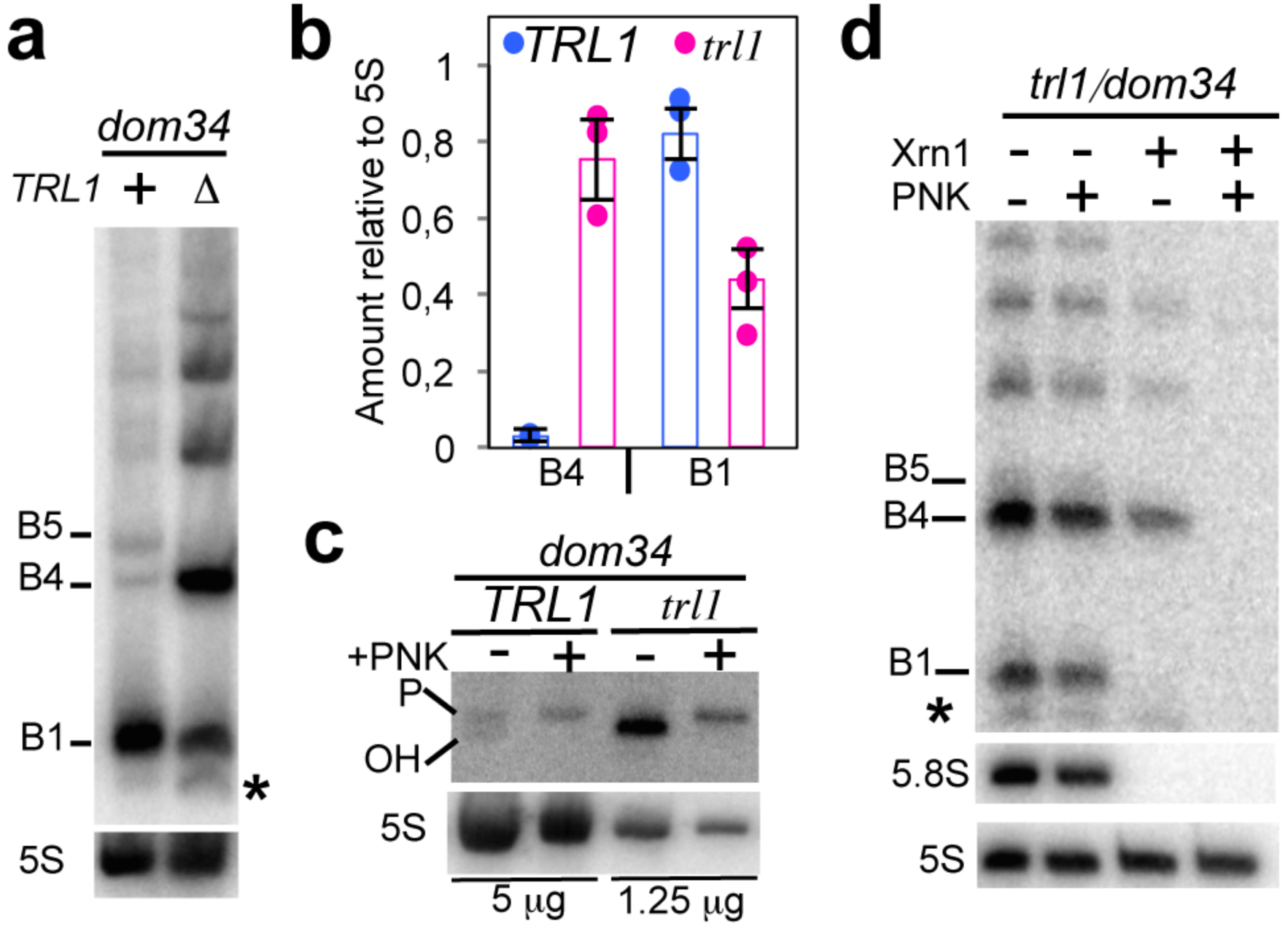
Endonucleolytically cleaved 5’-OH RNAs are phosphorylated by Trl1. **a** 8% PAGE followed by northern blot analysis using probe prA. Levels of 3’-NGD RNA fragments in *trl1/dom34* cells compared with those from *TRL1/dom34* cells. **b** B1 and B4 RNA quantification relative to 5S rRNA from three independent experiments as shown in (**a**). **c** 12% PAGE followed by northern blot analysis using probe prA. Treatment using T4 PNK to determine 5’-OH and 5’-P B4 RNA positions in the indicated strains. One-fourth of *trl1/dom34* total RNA treated was loaded to limit scan saturation and allow *TRL1/dom34* B4 RNA detection. The 5S rRNA served as a loading control. **d** As in Fig. 3a, Xrn1 digestion of total RNA extracts from *trl1/dom34* mutant cells in the presence or absence of T4 PNK treatment *in vitro*. A minor band detected in *trl1* is indicated by an asterisk (see also Supplementary Fig. 5 in which this band is detectable in *TRL1* cells). Error bars indicate standard deviation (s.d) calculated from three independent experiments. Source data are provided as a Source Data file.

### NGD cleavage products derived from mRNAs containing rare codons

We asked whether we could identify endonucleolytic cleavages on another NGD-targeted mRNAs, using what we learned about this process on truncated mRNAs. We chose an mRNA containing four contiguous rare CGA codons, which we call (CGA)_4_-mRNA, as an NGD target (Fig. 6a and Supplementary Fig. 6a)^5^. Similar to the truncated mRNAs, ribosomes were shown to stall when decoding rare codons, producing 5’- and 3’-NGD RNAs (Fig. 6a). As previously demonstrated^5, 35^, we detected 3’-NGD RNAs fragments in *dom34* or *DOM34* genetic contexts by northern blotting experiments using probe prB (Supplementary Fig. 6b). The precise identification of endonucleolytic cleavages by primer extension experiments is known to be challenging^5^ probably because, in contrast to truncated mRNAs, the positioning of ribosomes on contiguous rare codons is variable. We first asked whether we could detect 5’-NGD RNAs (Fig. 6a) using the same procedure as for NGD-targeted truncated RNAs (Fig. 4a, b). By probing the (CGA)_4_-mRNA in a large region upstream of the four CGA codons (Supplementary Fig. 6a), we detected RNA bands using a probe annealing 71 nts upstream of the first rare codon (probe prH, Fig. 6b and Supplementary Fig. 6a). Similar to the 5’-NGD RNAs produced from NGD-targeted mRNA1 (Fig. 4b), RNA detection required a *dom34* genetic background (Fig. 6b). The profile of the 5’-NGD RNAs resulting from endonucleolytic cleavages of (CGA)_4_-mRNA was remarkably similar to the B1, B4 and B5 RNAs from the truncated mRNA1. We then treated these RNAs with Xrn1 and, as anticipated, we observed that the ∼71-nt RNA, like the B4 RNA, was Xrn1-resistant, and that ∼47-nt and ∼77-nt RNAs, like the B1 and B5 RNAs, were Xrn1-sensitive (Fig. 6c). These results strongly suggest that NGD-targeted (CGA)_4_-mRNAs are a similar source of truncated RNAs which are, in turn, processed like mRNA1 by the NGD pathway.

**Fig. 6.**
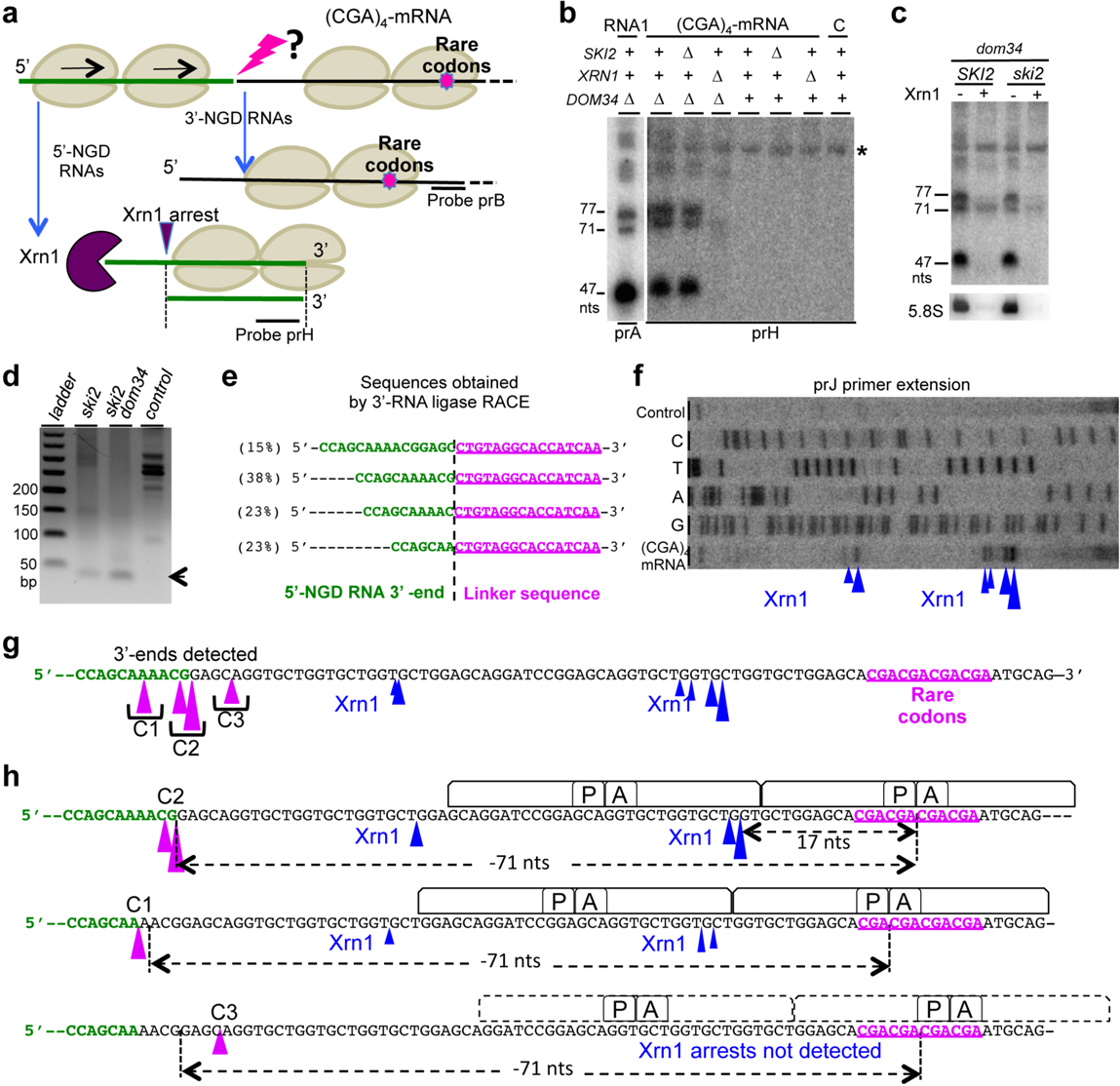
Identification of the endonucleolytic cuts on the NGD targeted (CGA)_4_-mRNA. **a** Schematic view of the (CGA)_4_-mRNA. 5’- and 3’-NGD RNAs are products of NGD. The lightning flash represents the potential endonucleolytic cleavage upstream of the ribosome stall site. Probes prB and prH are indicated. 5’-NDG RNAs are shown processed by Xrn1 as described in Fig. 4a, 4d and Supplementary Fig. 4. **b** Northern blot analysis using probe prH showing steady state levels of RNAs in the indicated mutant strains. Same membrane has been probed with prA as a ladder, and sizes of mRNA1 products such as B5 (77 nt), B4 (71 nt) and B1 (47 nt) are indicated. Only the *dom34* lane is shown. See Supplementary Fig. 6a for the sequence probed by prH. The 5S rRNA served as a loading control. Total RNA from WT cells without (CGA)_4_-mRNA expression served as a control, noted C. A non-specific band is indicated by an asterisk. **c** Xrn1 treatment *in vitro* of total RNA from *dom34* or *dom34/ski2* mutant cells and northern blot using probe prH. The 5.8S rRNA is a positive control of Xrn1 treatment. **d** PCR products obtained from 3’-RACE (see also Fig. 3c). Prior to PCR, cDNAs were produced from cells expressing (CGA)_4_-mRNA. Total RNA from cells without (CGA)_4_-mRNA expression served as a control. **e** Sequences obtained after 3’-RACE performed in (**d**) on *ski2*/*DOM34* total RNA. Sequence distribution is given in percentage. **f** Primer extension experiments using probe prJ to determine the 5’-end of RNAs. Xrn1-specific arrests are indicated by arrowheads. **g** Positioning of 3’-ends detected by 3’-RACE on (CGA)_4_-mRNA from *ski2/DOM34* cells (magenta arrowhead). Arrowhead sizes are proportional to the relative number of sequences obtained. Three cleavage clusters, C1, C2 and C3 were defined (see Supplementary Fig. 6d). Xrn1 arrests deduced from (**f**) are indicated by black arrowhead with sizes proportional to the intensity of reverse stops observed in (**f**). **h** Schematic view of the ribosome positioning on (CGA)_4_-mRNA deduced from Xrn1 arrests combined with the positioning of endonucleolytic cleavages provided by 3’-RACE. Source data are provided as a Source Data file.

The detection of short RNA species by prH probe suggested that endonucleolytic cleavages occurred just downstream, in a region located ∼70 nts upstream of the cluster of rare codons (Supplementary Fig. 6a). We thus set out to map the NGD cleavage sites on the (CGA)_4_-mRNA, using 3’-RACE for the detection of the 3’-ends of 5’-NGD RNAs in *ski2* and *ski2/dom34* mutant cells (Supplementary Fig. 6c). We obtained major RT-PCR products of about 45 bp that were purified, cloned and sequenced (Fig. 6d and Supplementary Fig. 6d). The 3’-end sequences (Fig. 6e) formed three clusters, C1, C2 and C3 (Fig. 6g), that map to ∼71 nts upstream of the second, third and fourth rare codon, respectively, consistent with cleavage within the footprint of the third ribosome as seen for the truncated mRNA1. No 3’-ends were detected within the region covered by two ribosomes, comforting the notion that disomes are not competent for NGD endonuclease activation. Xrn1 arrests mapping to 17-18 nts upstream of the A-site of the two first ribosomes positioned with either the second or third CGA codon in the A-site were also detected by primer extension assay (Fig. 6f-h). The strongest Xrn1 arrests corresponded to those where the lead ribosome contains the third CGA codon in the A-site (Fig. 6h), suggesting that the major stall occurs on this codon. Typically, Xrn1 is preferentially blocked 17 nts upstream of the first ribosomal A-site residue^26^. We speculate that this 1-nt difference reveals distinct conformations of stalled ribosomes on rare codons versus truncated mRNAs. All these results taken together suggest that the (CGA)_4_-mRNA and truncated mRNA1 are NGD-targeted in a highly similar process that results in cleavage within the footprint of the third ribosome, 71 nts upstream of the stall site for the leading ribosome.

Cleavages have been proposed by others to occur in the region covered by disomes using primer extension experiments^5^, ^29^. We thus analysed an mRNA containing rare codons with an identical ribosome stalling sequence to that previously examined^5^ (Supplementary Fig. 6e). We demonstrate that primer extension arrests detected in the region covered by disomes are abolished in *dxo1/xrn1* mutant cells, suggesting that they are the products of subsequent trimming by these enzymes. In conclusion, our data suggests that stalled disomes on truncated mRNAs or on mRNAs containing short CGA repeats are poorly competent for NGD endonucleolytic cleavages. Endonucleolytic cleavages instead occur upstream of collided disomes, in agreement with other 3’-RACE analyses^27^. Our data suggests that these cleavages first occur within the mRNA exit tunnel of the third stacked ribosome and those queuing further upstream.

## Discussion

In this study, we first characterized the 3’-NGD RNA fragments produced near the 3’-end of truncated mRNAs that mimic natural cleaved mRNAs known to be NGD targets. One advantage of studying the 3’-NGD products of truncated mRNAs is that the precise positioning of stalled ribosomes results in 3’-NGD RNA fragments of specific sizes. Indeed, the precise identification of endonucleolytic cleavages is known to be challenging for mRNAs containing rare codons^5, 29^ because the positioning of ribosomes on multiple contiguous rare codons is variable. Using a ribozyme to efficiently generate precise 3’-ends within an open reading frame, we were able to obtain detailed information about ribosome positioning on 3’-NGD RNAs, and provide the first precise mapping of the original site of endonucleolytic cleavage on an NGD substrate. Our model suggests that this cleavage occurs 71 nts from the 3’ end of the truncated mRNA, 8 nts upstream of the P-site codon in the third stacked ribosome (Figs 1g and 7). This localizes the 5’-extremity of cleaved RNA fragment within the mRNA exit tunnel, 4 nts downstream of the expected nucleotide position of a canonical mRNA that emerges from the ribosome and becomes available for cleavage by RNase I *in vitro*, classically used in ribosome foot-printing studies. A buried cleavage site is consistent with the idea that the NGD endonuclease might be the ribosome itself. However, we cannot fully exclude the possibility that the stalled ribosome allows access to an external nuclease with a specific conformation to penetrate this far into the mRNA exit tunnel.

**Fig. 7.**
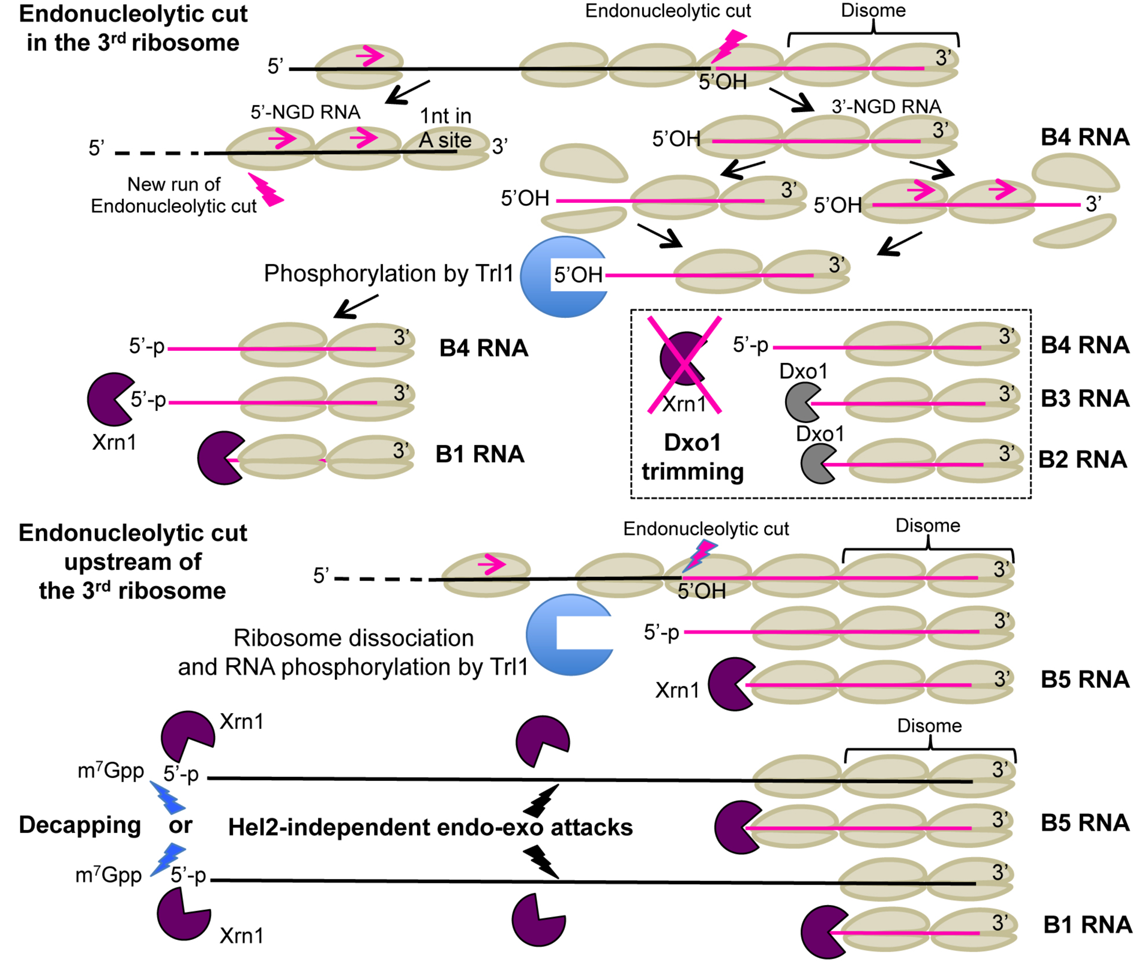
Model of No-Go decay pathway involving Trl1 kinase and 5’-3’ exoribonucleases. Top of figure, the third ribosome is represented as competent for NGD endonuclease activation. We propose that the two first stalled ribosomes are not properly conformed to trigger the endonucleolytic process. NGD endonuclease cleavage (lightning flash) occurs 8 nts upstream of the first P-site residue, within the mRNA exit tunnel of the ribosome. Upstream ribosomes covering the resulting 5’-NGD fragments can advance and stall on the new 3’-end with 1 nt in the ribosomal A-site. Colliding ribosomes on this new RNA fragment can induce a novel NGD endonuclease activation. On 3’-NGD RNAs, like B4 RNAs, the NGD-competent ribosome dissociates and facilitates access of Trl1 RNA kinase to the 5’-hydroxylated 3’-NGD RNA, but we cannot exclude that the leading ribosome dissociates and upstream ribosomes run to form a new disome with 5’-protruding RNA. Once the RNA is 5’-phosphorylated, the processive 5’-3’ exonucleolytic activity of Xrn1 can degrade and produce B1 RNA. Alternatively, upon Xrn1 inactivation, 5’-3’ exonucleolytic digestion of this RNA by Dxo1 can occur and produce trimmed RNAs, such as B3 and B2 RNAs. Middle of figure, upstream of the 3^rd^ ribosome, ribosomes are also competent for NGD endonuclease activation. Here, the endonucleolytic cleavage occurs in the 4^th^ ribosome and B5 RNAs can derive from such RNAs after phosphorylation and 5’-3’ digestion. Bottom of figure, alternative pathways are proposed: B1 and B5 RNA production could be initiated by decapping or *via* Hel2-independent uncharacterized endo/exonucleolytic attacks.

Xrn1 treatment of various mutant cell extracts suggested that the predominant ribosome configuration on truncated mRNAs is disomes. Interestingly, the existence of disomes on truncated mRNAs has been previously reported in ribosome profiling analysis^20^ and stacking of two or more ribosomes has been proposed as a prerequisite for the activation of the endonuclease^27^. The latter observation led to the proposition that ribosome collision triggers NGD cleavage upstream of disomes. Our data suggests that NGD endonucleolytic cleavage detected within the third stalled ribosome, suggests that the first two stalled ribosomes (disome) are not competent for the activation of the endonuclease activity (Fig. 7). Our 3’-RACE experiments did not amplify DNA products corresponding to RNAs matching the predicted sizes of NGD-cleaved RNAs within the second (41 nts) or first stalled ribosome (15 nts) (predicted sizes 95 and 125 nts, respectively), indicating that they do not occur to any significant level. The major ∼65-bp RT-PCR product corresponded perfectly to an RNA cleaved 71nt upstream of the 3’-extremity of mRNA1, suggesting this is the primary site of NGD cleavage. This also suggests that the particular conformation of disomes on these mRNAs is incompatible with an NGD endonuclease activity cutting upstream of the ribosome P-site. The ability to induce this precise NGD cleavage appears thus to be a normal property of stalled ribosomes, with disomes (and monosomes) being exceptions.

We considered the possibility that the *dom34* mutation may exaggerate the ribosome stalling and allow cleavage beyond what would naturally be observed. As discussed in the introduction, the analysis of NGD RNA fragments is facilitated by the *dom34* mutation and is crucial for RNA stabilization when analysing truncated mRNAs by northern blotting experiments^5, 20^. In the presence of Dom34, and more efficient ribosome dissociation, the exosome would certainly be more actively involved once the first endonucleolytic cleavage event has occurred^5, 20^. Importantly, however, our 3’-RACE experiments confirmed the existence of 5’-NGD products with 3’-ends matching the 5’-extremity of the 3’-NGD B4 RNA in cells containing Dom34 (Fig. 3c, d). Thus, NGD endonucleolytic cleavage does not occur randomly upstream of the ribosomal stall site and is not an artefact of *dom34* genetic context.

These observations were used to map endonucleolytic cleavages that occur on a second NGD-target mRNA containing rare codons, also in a *DOM34* genetic context. Endonucleolytic cleavages occurred 71nts upstream of the first residue in the leading ribosome A-site, in the region potentially covered by the third ribosome. Therefore, mRNAs containing rare codons are processed similar to truncated RNAs, although cleavage accuracy is slightly affected, correlating with specific Xrn1 arrests (Fig. 6f) that might be explained by a particular conformation of the first stalled ribosome^29^.

It has been recently reported that Cue2 is the endonuclease that cleaves NGD/NSD targeted RNAs^36, 37^. Its action has been proposed to occur in the ribosome A site and therefore deviates significantly from our observations. We do not have a good explanation for this difference. However, previous experiments with the naturally occurring truncated HAC1 mRNA are consistent with our mapping of the endonucleolytic cleavage site^20^. The HAC1 intron is known to be excised by Ire1, but RNA ligation can be incomplete and lead to a truncated but translated mRNA^38, 39^. Green and colleagues showed by ribosome profiling analysis that ribosomes stall at the 3’-end of the first exon of the HAC1 mRNA, leading to an endonucleolytic cleavage ∼70 nt upstream from the 3’-end^9^. Consistent with this observation, our data suggests that the NGD endonuclease cleaves 71 nt upstream of the 3’-end of the truncated mRNA1 (B4 RNAs, Fig. 1g). We also showed that the NGD endonuclease cleaves mRNA1 in the +1 reading frame. As a consequence, upstream ribosomes run forward on the 5’-NGD mRNA and stall with 1 nt in the A-site (Fig. 1g and 7). This is in agreement with ribosome profiling analysis reporting the predominance of short RNAs having one 3’-nucleotide in the ribosome A site^20, 36^.

We showed that the Trl1 kinase plays a role in NGD, and we propose that the resistance of the B4 3’-NGD RNA fragments to Xrn1 attacks *in vitro* and *in vivo* (Fig. 3a and Supplementary Figs 1b and 3a) is a direct consequence of the 3^rd^ ribosome preventing access to Trl1, with dissociation of this ribosome triggering the 5’-phosphorylation of the B4 RNA (Fig. 7). It is also possible that the leading ribosome dissociates, and that upstream ribosomes run forward to form a new disome with a 5’-protruding RNA (Fig. 7). Once the RNA is 5’-phosphorylated, the processive 5’-3’ exonucleolytic activity of Xrn1 can degrade to produce the disome protected B1 RNA (Fig. 7). The fact that RNAs longer than B5, which accumulate in the *trl1* mutant (Fig. 5a), are Xrn1-resistant (Fig. 5c and Supplementary Fig. 5) coupled with the absence of the B5 RNA upon Xrn1 inactivation (Fig. 1b-d) lead us to propose that the B5 RNA originates from 5’-OH RNAs cleaved within ribosomes upstream of the 3^rd^ ribosome, are phosphorylated by Trl1 and are digested by Xrn1 until it bumps into downstream ribosomes (Fig. 7). A portion of the B1 species (Fig. 5a and Supplementary Fig. 5a) is detected upon Trl1 inactivation, however. Therefore, 5’-phosphorylation of 5’-hydroxylated B4 RNAs only accounts for a portion of B1 RNA production. In agreement, the *hel2* mutation abolishes B4 RNA detection without impacting B1 and B5 RNA levels (Supplementary Fig. 3b). Mutations can thus reveal the existence of alternative pathways. In this regard, we learned that the inactivation of Xrn1 can lead to a Dxo1 trimming and production of B2 and B3 RNAs (Fig. 7). We cannot exclude the possibility that a portion of the B1 and B5 RNAs derive from alternative pathways such as the canonical 5’-3’ decay pathway (*i.e*, decapping and Xrn1 5’-3’ digestion)^24^ or an uncharacterized endo and/or exonucleolytic decay pathway (Fig. 7). Lower levels of NGD RNAs in a *dcp2* mutant (Fig. 1b, c) do not contradict a decapping requirement in some of their production, but the poor growth of *dcp2* mutant cells prevents us from drawing further conclusions. The potential existence of alternative pathways does not overshadow the strong impact of the *trl1* mutant on NGD RNA production (Fig. 5).

We show that the NGD pathway produces RNAs bearing a 5’-hydroxyl group (Figs 3a, 5 and 7). 5’-hydroxyl and 2’-3’ cyclic phosphate 3’ extremities are typically generated by metal-independent endoribonucleolytic reactions^40^. We also show that the Trl1 kinase phosphorylates these 3’-NGD fragments to allow degradation by 5’-3’ exoribonucleases. Thus, in addition to its role in tRNA splicing and in phosphorylation of 5’-hydroxylated exon of the HAC1 mRNA^34^, Trl1 is an important player in NGD pathway. This study also provides mechanistic insights that will help to go further in the comprehension of mRNA surveillance pathways in connection to NGD.

## Methods

### Yeast Media, plasmids, strains, and oligonucleotides

The media, strains of *S. cerevisiae,* oligonucleotides, ***s*ynthetized DNA*s*** and plasmids used in this study are described in the Supplementary Methods and Supplementary Tables 1, 2, 3 and 4 respectively.

### Northern blot analysis

8 OD_600nm_ of exponentially growing cells were collected by centrifugation. RNA was extracted by adding on cell pellet 500 µl of buffer AE (50 mM Na acetate [pH 5.3], 10 mM EDTA), vortexing, and adding 500 µl of phenol previously equilibrated in buffer AE. After vortexing, the tubes were incubated at 65°C for 4 min and then frozen in a dry ice-ethanol mix, followed by a 15-min centrifugation at room temperature. A second extraction with 500 µl of phenol-Tris-EDTA-chloroform was performed. RNA from 360 µl of supernatant was precipitated with 40 µl of 3 M Na acetate (pH 5.3) and 2.5 volumes of ethanol and centrifuged at 4°C for 30 min. The pellet was washed in 80% ethanol, dried at room temperature and resuspended in water. After electrophoresis, RNA was blotted onto a Hybond-N+ membrane (Amersham) and hybridization to end-labeled oligodeoxynucleotide probes was carried out at 42°C overnight in Roti-Quick solution (Carlroth). The Hybond-N+ membrane was washed successively at 42°C in 5× SSC–0.1% SDS and twice in 1× SSC–0.1% SDS. Blots were exposed to PhosphorImager screens, scanned using a Typhoon FLA 9500 (Fuji), and quantified with ImageJ software.

### Gel electrophoresis for separation of RNA molecules

Total RNA was resolved by 8% TBE-Urea polyacrylamide or 1.4% TBE-Agarose gels, and to dissociate 5’-hydroxylated from 5’-phosphorylated RNAs, total RNA was resolved by 12% TBE-Urea polyacrylamide gels. All gels were followed by northern blot analysis.

### *In vitro* RNA digestion

RNA digestion of 20 OD_260nm_ of cell extracts were performed by using 1 unit of Xrn1 (Biolabs) in NEB buffer 3 at 25°C during 30 min unless otherwise indicated. NEB Buffer 3 was replaced by T4 PNK buffer (NEB) in kinase assays in the presence or absence of Xrn1 (Figs 3a and 5d). For RNase I treatment of cell extracts, 20 OD_260nm_ of extracts (prepared without heparin) were incubated with 0.5, 1, or 2 µl of RNase I (Invitrogen, 100 units/µl) 30 min at 25°C. For total RNA treatment, 5 µg of RNA were digested 30 min at 25°C. All RNase treatments were followed by RNA extraction and northern blot analysis as described above.

### Polysome Analysis

Yeast cells were grown exponentially to 0.8 OD_600_ at 28°C and the 200 ml cell culture was quickly cooled in ice water and then centrifuged and washed in 20 ml of buffer A containing 10 mM Tris pH 7.4, 100 mM NaCl and 30 mM MgCl_2_, and 0.5 mg/ml heparin. Heparin was only omitted for RNase I treatment^24^. Cycloheximide was not used to prevent any drug–induced ribosome positioning. The cell pellet was resuspended in 0.6 ml of buffer A. A 1.4 g amount of glass beads was added, and cell lysis was performed by bead beating twice for 30 secondes each time (with intermittent cooling). After lysis, the cell extract was clarified for 5 min at full speed in a microcentrifuge. The equivalent of 20 OD_260nm_ units of cell extract (80 μg of total RNA) was then layered onto linear 10% to 50% sucrose density gradients. Sucrose gradients (in 10 mM Tris-HCl [pH 7.4], 70 mM ammonium acetate, 30 mM MgCl_2_) were prepared in 12 × 89 mm polyallomer tubes (Beckman Coulter). Polysome profiles were generated by continuous absorbance measurement at 254 nm using the Isco fraction collector. Gradients were collected in 20 fractions and processed for northern blotting as described above.

### Primer Extension

Radiolabeled primers (primers prA and prE for mRNA1, and primer prJ for (CGA)_4_-mRNA) were used and Maxima H Minus reverse transcriptase (ThermoFisher) was used to synthesize a single-stranded DNA toward the 5’-end of the RNA. The size of the labeled single-stranded DNA was determined relative to a sequencing ladder (ThermoFisher Sequenase sequencing kit) on 5% TBE-Urea polyacrylamide gel. Oligonucleotides were radio-labeled with [γ-32P]ATP with the T4 polynucleotide kinase (NEB).

### 3’-end RNA mapping

Mapping was performed according to the 3’-RNA ligase mediated RACE method^31^ with minor modifications: Total RNA preparations were first 3’-dephosphorylated using T4 PNK for 1h at 37°C without ATP and pre-adenylated linker (Universal miRNA cloning linker, NEB) ligation was performed during 4h at 22°C in the presence of truncated RNA ligase 2 (NEB)^30^. Reverse transcriptase reactions were performed using primer prE complementary to the linker sequence. PCR primer prF specific to mRNA1, or primer prK specific to (CGA)_4_-mRNA, were used with primer prE in PCR reactions (Supplementary Figs 3a and 6c). PCR products were purified, cloned into Zero Blunt TOPO PCR Cloning vector (Invitrogen), transformed and plasmids sequenced.

### Data availability

Uncropped and unprocessed scans of all blots and all calculations are supplied in the Source Data file at FigShare with DOI information (0.6084/m9.figshare.11317613)

### Data availability statement

The authors declare that all data supporting the findings of this study are available within the article and its supplementary information files or from the corresponding author upon reasonable request. The source data underlying Figs 1b-f, 2a-b, 3a, c, 4b, c, 5a-d and 6b-d, f and Supplementary Figs 1b, d-i, 2a-c, 3a, b, 5 and 6b, e are provided as a Source Data file in Supplementary information and at FigShare with DOI information (0.6084/m9.figshare.11317613).

## End Notes

## Author contributions

A.N., S.C., R.S.F., J.H., C.T. and L.B. designed, performed and analysed data. L.B. wrote the manuscript. A.N. and S.C. contributed equally and are listed in chronological order.

## Competing Interests

The authors declare no competing interests.

## Acknowledgements

This work has been supported by AAP Emergence Sorbonne Université, SU-16-R-EMR-03 and by the “Initiative d’Excellence” program from the French State grant “DYNAMO”, ANR-11-LABX-0011-01. S.C. was a recipient of fellowship from the Ministère pour la Recherche et la Technologie (MNRT). A.N. was supported by a doctoral grant from “DYNAMO”, ANR-11-LABX-0011-01. We thank Jay Hesselberth for providing *trl1* mutant and Jeff Coller for the *dcp2Δ* mutant. We thank Ciaran Condon, Josette Banroques and Kyle Tanner for technical assistance. We thank Ciaran Condon for comments on the manuscript and constructive discussions.

## Competing financial interests statement

The authors declare no competing financial interests.

## Supplementary file

File name; Supplementary Information

Description: this pdf file contains 2 Supplementary Notes, 6 Supplementary Figures and 4 Supplementary Tables.

File name: Source Data File

Description: xls file containing all uncropped and unprocessed scans underlying Figs 1b-f, 2a-b, 3a, c, 4b, c, 5a-d and 6b-d, f and Supplementary Figs 1b, d-i, 2a-c, 3a, b, 5 and 6b, e and all calculations corresponding to Figures 1f and 5b.

## Supplementary Information

### Supplementary Notes

**Supplementary Note 1** Ribosome protection of 3’-NGD RNA fragments from Xrn1 activity. We first focused on the fate of the major B1 (47 nts) and B5 (77 nts) RNA species detected in *dom34* cell extracts, which likely correspond to RNAs protected from Xrn1 digestion *in vivo* by trisomes and disomes, respectively. We showed that Xrn1 treatment of RNA in *dom34* cell extracts had no impact on B5 and B1 RNAs *in vitro,* suggesting that these RNA species are indeed protected by ribosomes (Supplementary Fig. 1b). Interestingly, the persistence of the 71-nt B4 RNA after Xrn1 treatment suggests that this RNA may also be protected by up to three ribosomes in the *dom34* background (Supplementary Fig. 1b). We also added purified Xrn1 to cell extracts of the *dom34/xrn1* strain *in vitro* and showed that it can efficiently recapitulate the production of the B1 species observed in Xrn1-containing cells *in vivo*. The appearance of the B1 RNA was inversely correlated to the amount of B4, B3 and B2 RNAs remaining, suggesting that these three species have unprotected 5’-protruding RNA extremities *in vivo,* due to the absence of the 5’-3’ exoribonuclease Xrn1 (Supplementary Fig. 1b). The B5 RNA appears to be also generated at the expense of some larger species by Xrn1 treatment *in vitro*, consistent with the presence of trisomes on this species in the *dom34/xrn1* cell extracts (Supplementary Fig. 1b). Based on these experiments, we propose that the B1 (47 nts) and B5 (77 nts) species correspond to Xrn1-trimmed RNAs protected by two and three ribosomes, respectively^1^ and that at least some portion of the 71 nt B4 RNA is also protected from Xrn1 by three ribosomes.

**Supplementary Note 2** Ribosome protection of 3’-NGD RNA fragments from RNase I activity. To validate the presence and number of ribosomes on 3’-NGD RNAs by a third method, and particularly the presence of trisomes on the 77-nt B5 and 71-nt B4 species, we also performed RNase I protection assays on cell extracts of *dom34* and *dom34 /xrn1* strains. We hypothesized that B5 and B4 RNAs protected from RNase I by three ribosomes should be detectable with both probes prA and prD (Supplementary Fig. 1c). The presence of two or three ribosomes on the major RNA species B1, B4 and B5 in *dom34* cells (deduced from primer extension experiments, ribosome association and Xrn1 treatment *in vitro*) would preferentially conduct RNase I to cleave at three major sites, Cut1, 2 and 3 in Supplementary Fig. 1c. After RNase I treatment (Supplementary Fig. 1d, e), the accumulation of RNase I protected RNAs of similar size to B5 is consistent with the hypothesis that this RNA is covered by trisomes in *dom34* mutant extracts (Fig. 1f). It is known that RNase I and Xrn1 cleave about 15 nts^2^ and 17 nts^1^ upstream of the first nt of the ribosomal A-site, respectively. We thus expected 5’-end of the B5 RNA to be 1-2 nts shorter than the Xrn1 product after RNase I treatment, if we had accurately calculated the positions of ribosomes on this RNA. Indeed, primer extension experiments confirmed that ribosomes protect a 75-nt species from RNase I in the *dom34* background (Supplementary Fig. 1f). The equivalent of the B1 RNA was 46 nt in size after RNase I treatment (Supplementary Fig. 1g). B4 RNA was not detected using probe prA (Supplementary Fig. 1d, h). We wondered whether RNase I cleaved preferentially at Cut3 (Supplementary Fig. 1c), thus preventing the detection of B4 using probe prA. We probed the membrane in Supplementary Fig. 1h with prA (Fig. 1i), and two distinct RNA species were detected corresponding to B5 and B4 processed by RNase I at Cut3, and inefficiently cleaved at the Cut2 site (Supplementary Fig. 1c). We thus propose that the 5’-extremities of B4 RNA are also protected by ribosomes (at least two ribosomes limiting RNase I attack at Cut2). We conducted the same experiment on the B4 RNA from *xrn1/dom34* cell extracts. These RNAs were sensitive to Xrn1 treatment *in vitro* (Supplementary Fig. 1b), and using probe prD to detect the B4 RNA specifically, we observed that these RNAs were also protected from RNase I to a similar extent as B4 in *dom34* cell extracts (Supplementary Fig. 1i). Thus, whether two or three ribosomes dwell on the 71-nt B4 RNA in *xrn1/dom34* mutant cell extracts is not completely clear as this species is sensitive to Xrn1 *in vitro* (*i.e.* have 5’-ribosome-free extensions that can be pared down to B1 by Xrn1 digestions) (Supplementary Fig. 1b), which would be consistent with protection by two ribosomes, but it is resistant to RNase I (Supplementary Fig. 1i), which is more consistent with three.

## Supplementary Methods

**Yeast Media.** Strains were grown in YPD medium or in synthetic minimum media (SD)^4^. Minimal media was completed for auxotrophy, leucine, histidine and/or uracil were omitted to keep selection for plasmids when necessary. 200 µg/ml G418 Sulfate (Geniticin, American Bioanalytical), 100 μg/ml Hygromycin B (Sigma-Aldrich) and 100 μg/ml ClonNat (Werner Bioagents) were added in YPD media plates to select for KanMX4, HphMX4 and NatMX6 respectively.

**Strains used in this study.** Mutant strains were generated by the one-step gene replacement using PCR fragment of the NatMX6 cassette amplified from plasmid pFA6a-natMX6^5^,with HphMX4 containing cassette amplified from pAG32^6^ or by the KanMX6 cassette amplified by PCR from plasmid pFA6a-kanMX6 respectively. Correct integration was confirmed by PCR with primers. See Supplementary Table 1 for strains, and Supplementary Table 2 for used primers.

**Plasmids used in this study.** Yeast plasmids used in this study were constructed using standard molecular biology procedures. To construct pLB138 (mRNA1RZ) and pLB127 ((CGA)_4_-mRNA), p415ADH1^7^ was first digested by SpeI-XhoI. DNA fragments containing URA3 were amplified by PCR from pRS316^8^ using primers olb592-olb593 and digested by SpeI-BamHI. In parallel, oligonucleotides olb-ins1-f and olb-ins1-r were annealed. All DNA fragments were ligated to build pADH1-URA3. pADH1-URA3 was then digested by BspEI-NdeI. Genomic DNA was amplified using primers olb594-olb596 and digested by BamHI-NdeI in order to insert an additional ORF (ORF2) in the 3’-region of *URA3*. In parallel, in order to insert a ribozyme sequence (Rz) just downstream URA3 sequence, oligonucleotides olb625 and olb626 were annealed and all DNA fragments were ligated to form pADH1-URA3-Rz-ORF2. To insert 4 CGA codons, oligonucleotides olb640 and olb641 were annealed and all DNA fragments were ligated to form pADH1-URA3-(CGA)_4_-ORF2. Additionally, oligonucleotides olb-2HA-f and olb-2HA-r were annealed and cloned into pADH1-URA3-Rz-ORF2 or pADH1-URA3-(CGA)4-ORF2 (XbaI-SpeI digestion). The resulting plasmids p415ADH1-2HA-URA3-Rz-ORF2 and p415ADH1-2HA-URA3-(CGA)_4_-ORF2 were named p138 and p127 respectively. The resulting ORF sequence of the mRNA with 3’-Rz insertion is shown in Supplementary Fig. 1a. Plasmids pDxo1_WT_ and pDxo1_mut_ used for the expression *in vivo* of WT Dxo1-Flag or of a catalytic mutant of Dxo1 (E260A D262A) were both created using synthetized DNAs (Genecust) cloned in SalI-XbaI sites of pRS313 (synthetized DNA sequences in Table S4). Thermocompetent NEB 10-beta *E. coli* (NEB) were used for cloning; all the plasmids were verified by sequencing (Eurofins Genomics).

**Supplementary Fig. 1.**
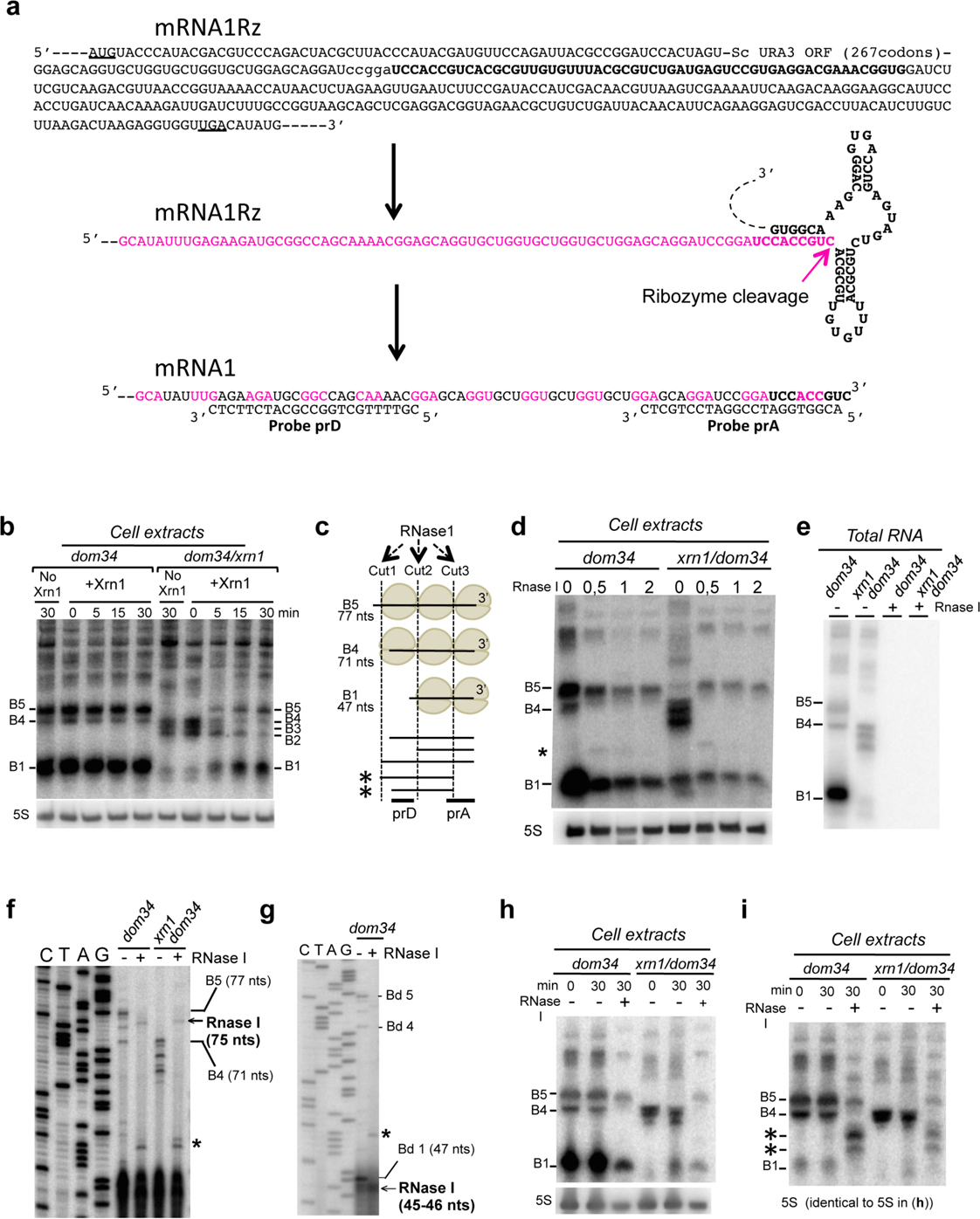
3’-NGD RNA fragment analysis, related to Fig. 1. **a** Sequence of mRNA1Rz. The translational start (AUG) and stop codon (UGA) are underlined. The ribozyme sequence is shown in bold. The magenta arrow indicates the ribozyme cleavage site. mRNA1 is the truncated stop-less codon mRNA after ribozyme cleavage. Probes prA and prD are indicated. **b** Xrn1 treatment *in vitro* of cell extracts (*i.e.* mRNAs in presence of ribosomes) from *dom34* or mutant cells, followed by RNA extraction and northern blot using probe prA. Sizes in nts are deduced from experiments shown in (Fig. 1c-d). **c** Schematic view of ribosomes covering RNA species B1, B4 and B5 observed in *dom34* mutant cells. Cut1, Cut2 and Cut3 represent potential RNase I cleavage sites. Probes prA and prD used in northern blots analysis shown in (**h**) and (**i**) are indicated. 5’-extremities of B5 and B4 RNAs potentially protected by two ribosomes and detected by prD are indicated by asterisks. **d** RNase I treatment *in vitro* of cell extracts (*i.e.* mRNAs in presence of ribosomes) from *dom34* or *dom34/xrn1* mutant cells followed northern blot using probe prA. The 5S rRNA served as a loading control. 0.5, 1, or 2 µl of RNase I were used (100 units/µl). **e** Similar RNase I treatment analysis that in (**d**) but on extracted RNAs (*i.e.* mRNAs in absence of ribosomes). **f**-**g** Primer extension experiments using probe prA to determine the 5’-end of RNAs after RNase I treatment of *dom34* and *xnr1/dom34* cell extracts as performed in (**d**). The band indicated by an asterisk is lost at a higher concentration of RNase I as shown in (**d**). **h** RNase I treatment of cell extracts *in vitro*, analysed as in (**b**). **i** The same membrane in (**h**) has been probed with prD. Source data are provided as a Source Data file.

**Supplementary Fig. 2.**
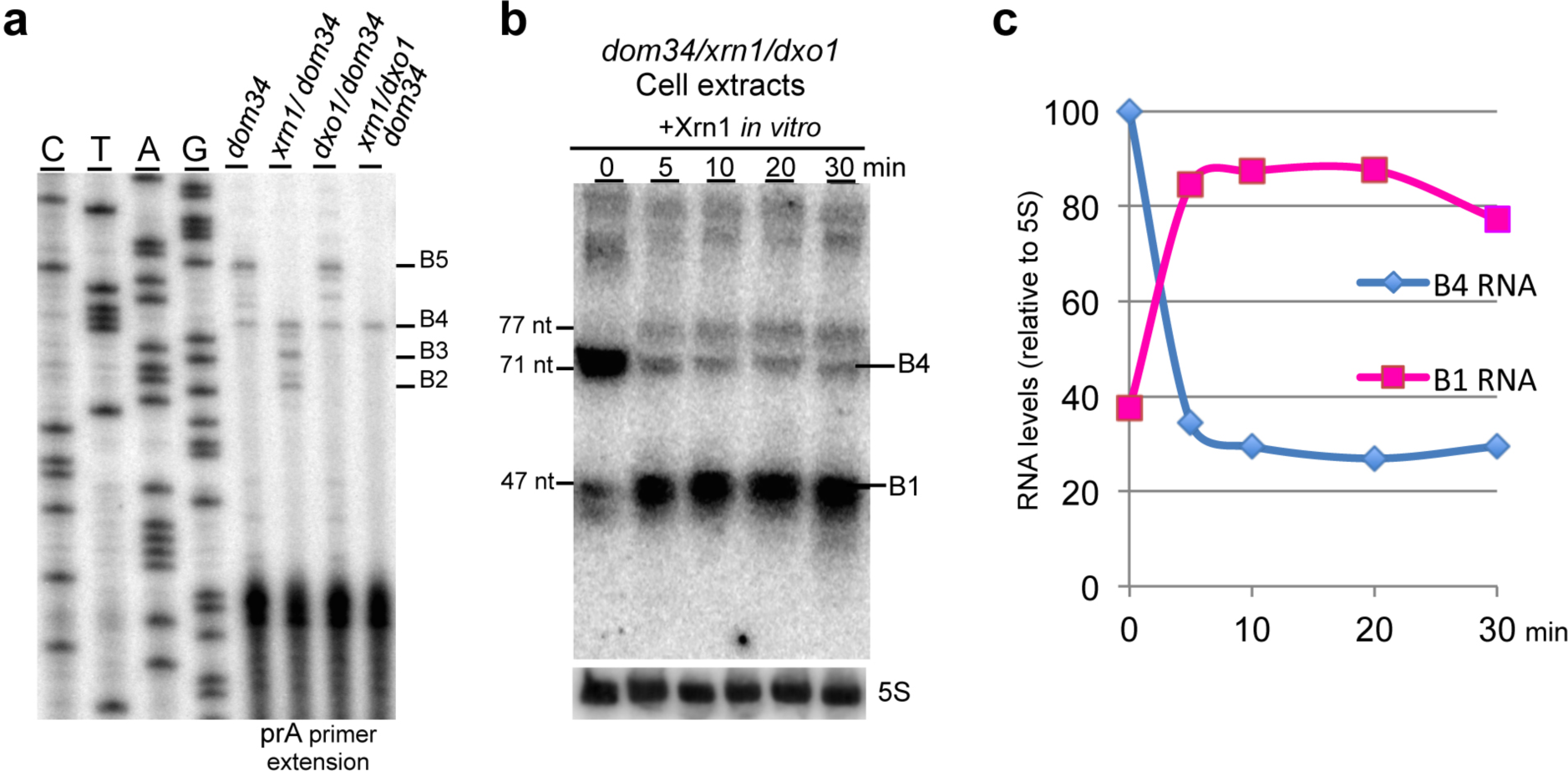
Dxo1 produces heterogeneity of 3’-NGD RNA fragments in Xrn1 deficient cells, related to Fig. 2. **a** Primer extension experiments using probe prA for determining the 5’-end of B1, B2, B3, B4 and B5 3’-NGD RNAs in indicated strains. **b** Cell extracts from *xrn1/dxo1/dom34* cells are digested by Xrn1 *in vitro*. 8% PAGE followed by northern blotting analysis using probe prA. **c** Quantification (a mean of two independent experiments) of RNAs shown in (**b**). B1 RNA (as the 47 nt species) and B4 RNA (as the 71 nt species) are indicated in amount relative to the 5S rRNA. Source data are provided as a Source Data file.

**Supplementary Fig. 3.**
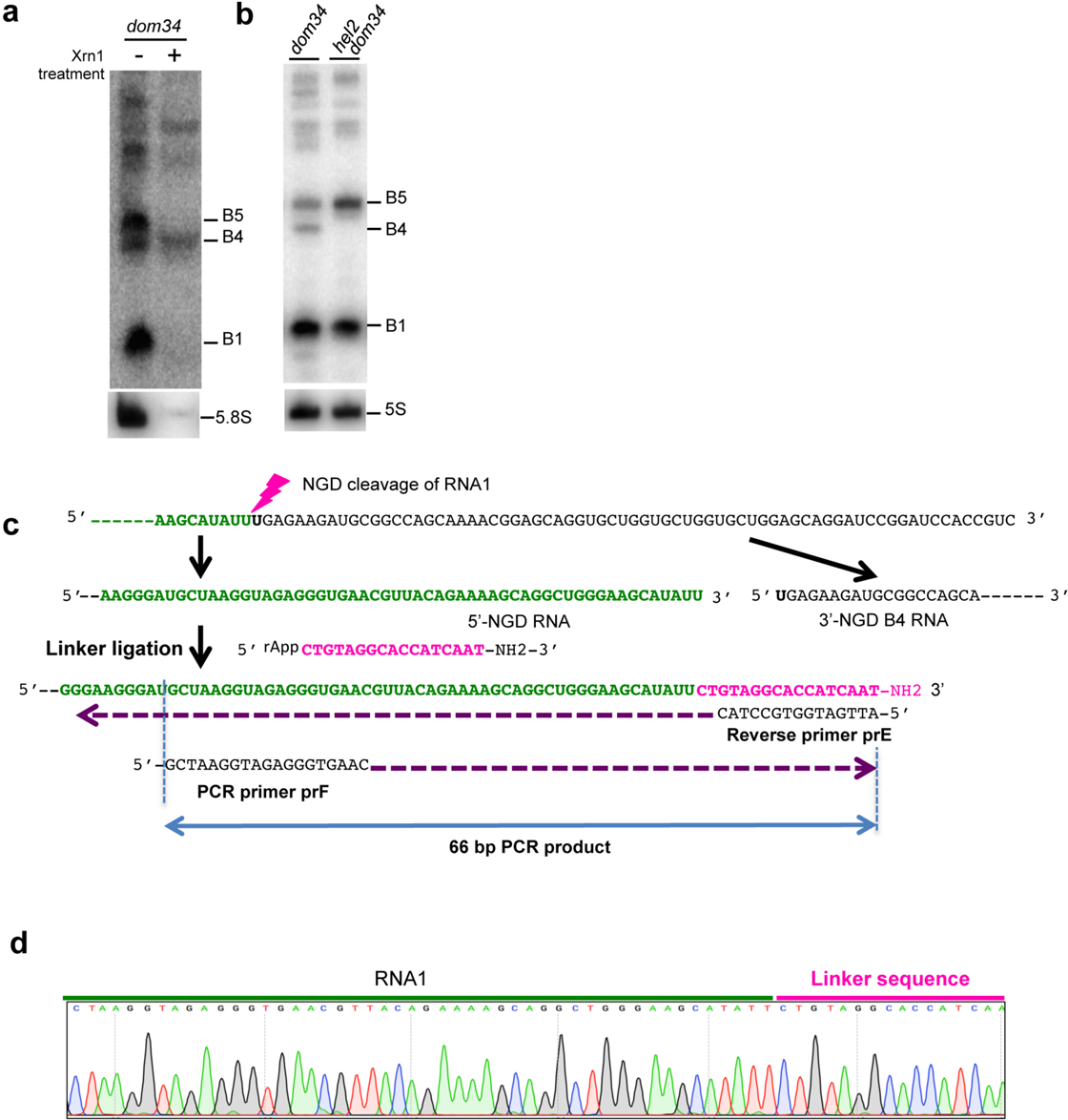
Characterization of B4 RNAs and 3’-RNA ligase mediated RACE, related to Fig. 3. **a** Xrn1 digestion of total RNA extracts from *dom34* mutant cells in Xrn1 buffer *in vitro*. 8% PAGE followed by northern blot analysis using probe prA. The 5.8S rRNA served as a positive control of Xrn1 treatment. **b** B4 RNA production is not detected in *hel2/dom34* mutant cells. 8% PAGE followed by northern blotting analysis using probe prA. **c** mRNA1 before and after the endonucleolytic cleavage (represented by the lightning flash) producing the 3’-NGD B4 RNA, and the resulting 5’-NGD RNA. The expected 3’-extremity is shown ligated to the universal miRNA linker (NEB). Sequence of reverse primer prE and PCR primer prF are indicated. A PCR product of 66bp is expected. **d** Chromatogram representing sequences obtained from 3’-RACE experiments performed on total RNA from *ski2* and *ski2*/*dom34* mutant cells. Source data are provided as a Source Data file.

**Supplementary Fig. 4.**
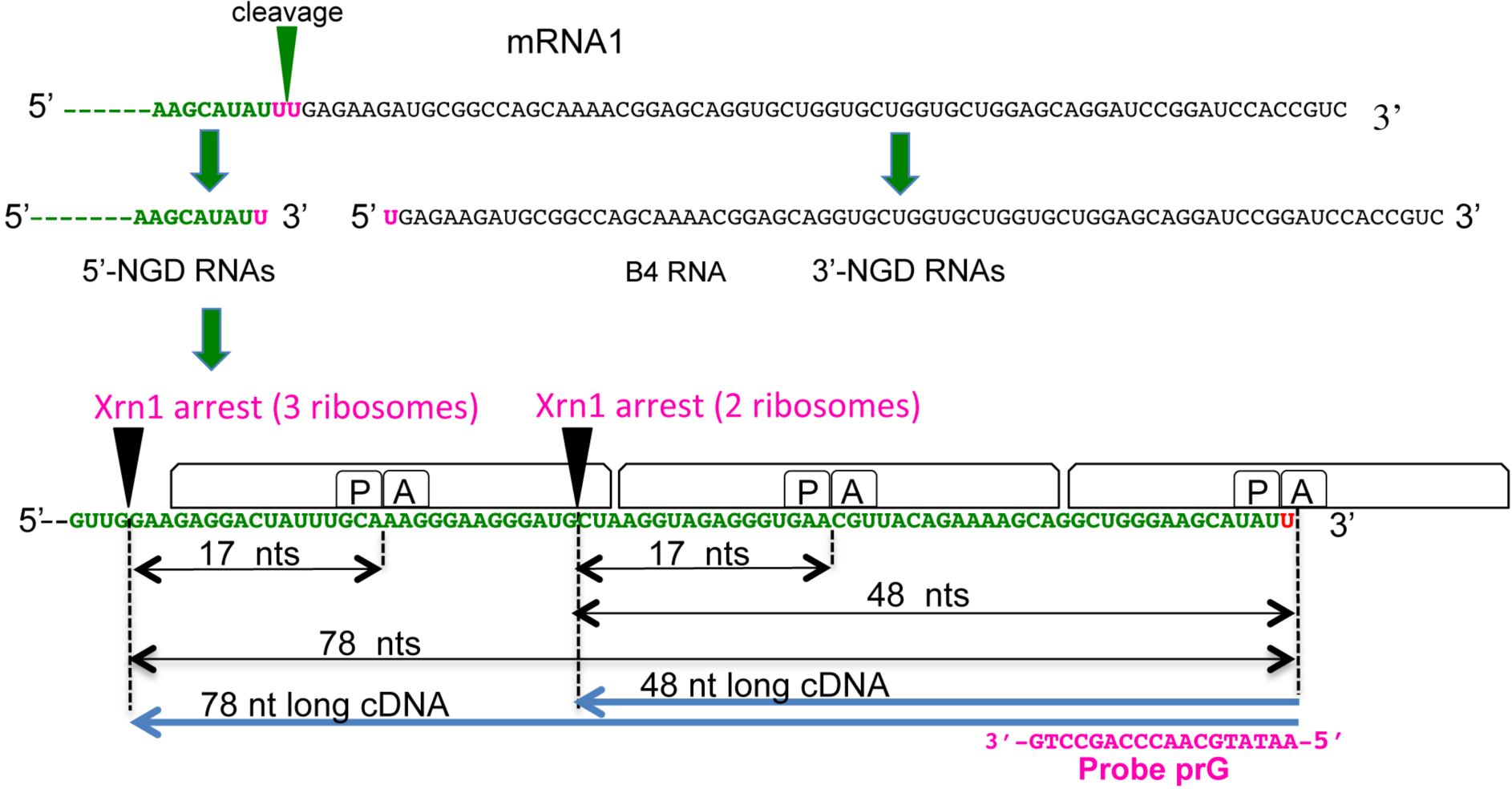
Analysis of the fate of 5’-NGD RNA, related to Fig. 4. Schematic model of mRNA1 before and after the endonucleolytic cleavage producing B4 RNA. The 5’-NGD resulting RNA is shown covered by ribosomes and is shown processed by Xrn1 in 48- and 78-nt RNAs when covered by two and three ribosomes, respectively. Xrn1 arrests occur 17 nts upstream of ribosomal A-site first residues^1^. Using probe prG in primer extension experiments, 48- and 78-nt cDNA products are expected.

**Supplementary Fig. 5.**
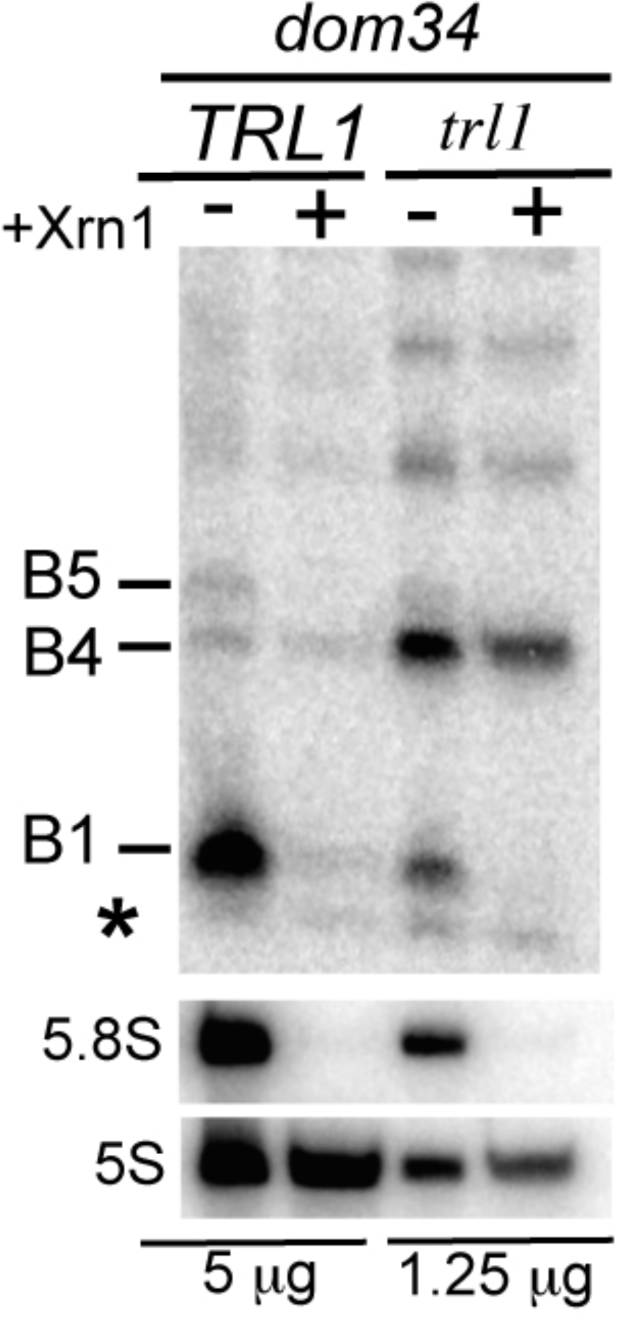
Related to Fig. 5, endonucleolytically cleaved 5’-OH RNAs are phosphorylated by Trl1. Xrn1 digestion of total RNA extracts from *TRL1/dom34* and *trl1/dom34* mutants in Xrn1 buffer. 5 μg and 1.25 μg of total RNA, from *TRL1* and *trl1* cell extracts respectively, were treated. A minor band detected in *TRL1* and *trl1* cells is indicated by an asterisk. The 5S rRNA served as a loading control. The 5.8S rRNA served as a positive control of Xrn1 treatment. Source data are provided as a Source Data file.

**Supplementary Fig. 6.**
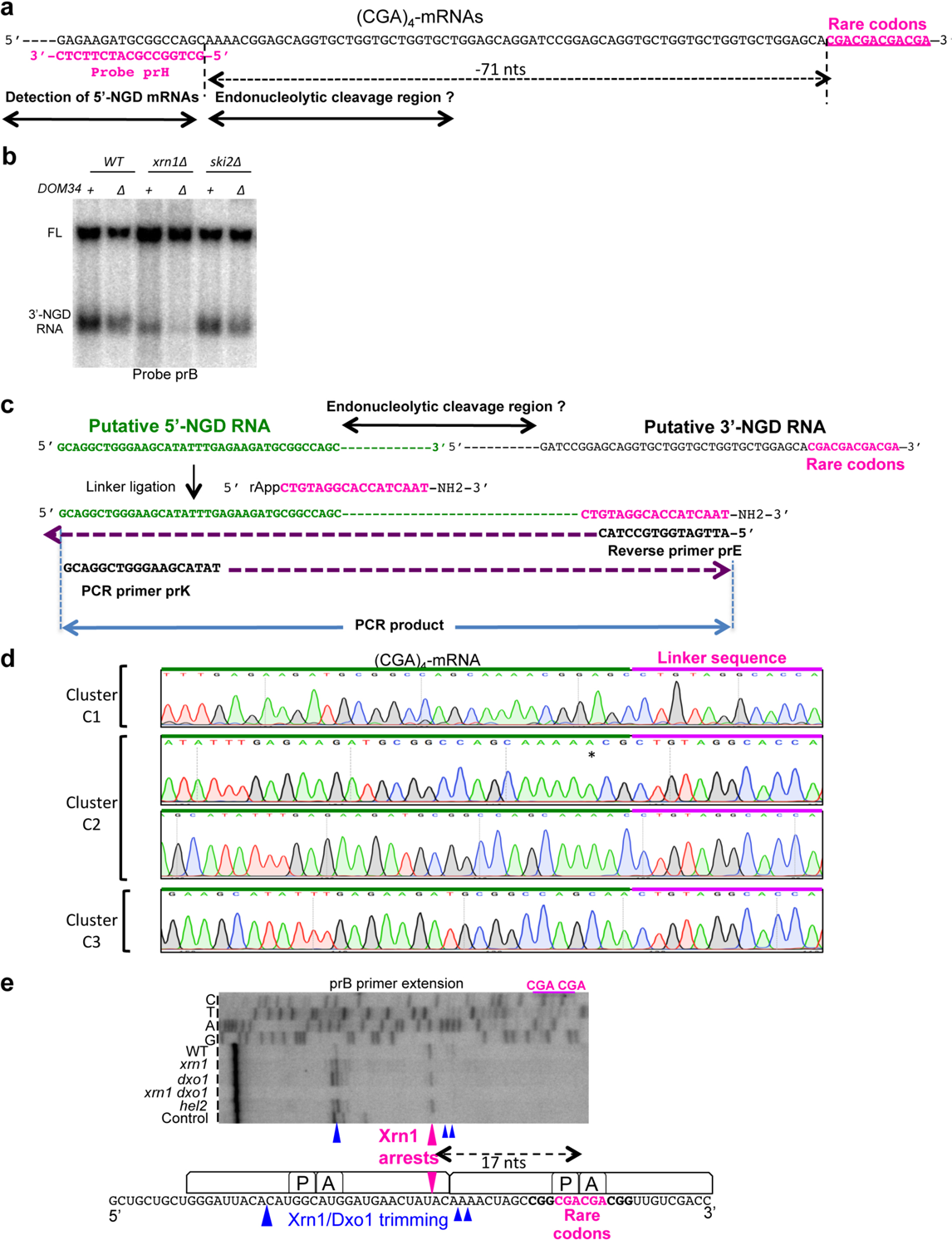
Analysis of (CGA)_4_-mRNAs, related to Fig. 6. **a** Partial sequence of (CGA)_4_-mRNA showing region upstream the four CGA rare codons. Positioning of probes prH is indicated. **b** 1.4% agarose gel followed by northern blotting analysis using probe prB showing steady state levels of RNAs in *dom34* and other indicated mutant strains. Full length (CGA)_4_-mRNA is noted FL, and the 3’-NGD RNAs are indicated. **c** 3’-RACE. The region of potential endonucleolytic cleavage, the 3’- and 5’-NGD RNAs are indicated. The putative 3’-extremity is shown ligated to the universal miRNA linker (NEB). Sequence of reverse primer prG and PCR primer prK are indicated. **d** Chromatogram representing sequences obtained from 3’-RACE experiments performed on total RNA from *ski2* mutant cells and the three cleavage clusters C1, C2 and C3. The asterisk indicates one nucleotide A mismatch found in sequences. **e** Primer extension experiments using probe prB to determine the 5’-end of the mRNA containing two contiguous CGA rare codons as described previously^3^. A schematic view of the ribosome positioning on this mRNA is shown below and Xrn1-specific arrest is indicated by a magenta arrowhead. Arrests dependent on Xrn1/Dxo1 activities are also indicated by blue arrowheads. Source data are provided as a Source Data file.

## Supplementary Tables

**Supplementary Table 1.** Strains used in this study

| Name | alias | genotype | ref. |  |
| --- | --- | --- | --- | --- |
| BY4741 | WT | *MATa his3Δ1 leu2Δ0 met15Δ0 ura3Δ0* | euroscarf |  |
| Y05329 | *dom34* | *MATa his3Δ1 leu2Δ0 met15Δ0 ura3Δ0 dom34::kanMX4* | euroscarf |  |
| BY11756 | *met22* | *MATa his3Δ1 leu2Δ0 lys2Δ0 ura3Δ0 met22::kanMX4* | euroscarf |  |
| Y04540 | *xrn1* | *MATa his3Δ1 leu2Δ0 met15Δ0 ura3Δ0 xrn1::kanMX4* | euroscarf |  |
| Y05307 | *ski2* | *MATa his3Δ1 leu2Δ0 met15Δ0 ura3Δ0 ski2::kanMX4* | euroscarf |  |
| YLB152 | *dxo1* | *MATa his3Δ1 leu2Δ0 met15Δ0 ura3Δ0 dxo1::kanMX4* | euroscarf |  |
| YLB177 | *met22 xrn1* | *MATa his3Δ1 leu2Δ0 lys2Δ0 ura3Δ0 met22::kanMX4 xrn1::hphMX4* | this study | derived from BY11756 |
| YLB082 | *ski2 dom34* | *MATa his3Δ1 leu2Δ0 lys2Δ0 ura3Δ0 ski2::kanMX4 dom34::natMX6* | this study | derived from Y05307 |
| YLB083 | *xrn1 dom34* | *MATa his3Δ1 leu2Δ0 lys2Δ0 ura3Δ0 xrn1::kanMX4 dom34::natMX6* | this study | derived from Y04540 |
| YLB084 | *met22 dom34* | *MATa his3Δ1 leu2Δ0 lys2Δ0 ura3Δ0 met22::kanMX4 dom34::natMX6* | this study | derived from BY11756 |
| YLB178 | *dxo1 dom34* | *MATa his3Δ1 leu2Δ0 met15Δ0 ura3Δ0 dxo1::kanMX4 dom34::natMX6* | this study | derived from YLB152 |
| YLB179 | *xrn1 dxo1 dom34* | *MATa his3Δ1 leu2Δ0 met15Δ0 ura3Δ0 dxo1::kanMX4 xrn1::hphMX4 dom34::natMX6* | this study | derived from YLB178 |
| YLB176 | *xrn1 met22 dom34* | *MATa his3Δ1 leu2Δ0 lys2Δ0 ura3Δ0 met22::kanMX4 dom34::natMX6 xrn1::hphMX4* | this study | derived from YLB177 |
| YLB302 | *ltn1 dom34* | *MATa his3Δ1 leu2Δ0 met15Δ0 ura3Δ0 ltn1::kanMX4 dom34::natMX6* | this study | derived from Y04540 |
| YLB303 | *hel2 dom34* | *MATa his3Δ1 leu2Δ0 met15Δ0 ura3Δ0 hel2::kanMX4 dom34::natMX6* | this study | derived from Y04540 |
| YJC432 | *dcp2* | *MATa his3Δ1 leu2Δ0 met15Δ0 ura3Δ0 dcp2::neoMX* | Jeff Collier | 9 |
| YLB156 | *dcp2 dom34* | *MATa his3Δ1 leu2Δ0 met15Δ0 ura3Δ0 dcp2::neoMX dom34::natMX6* | this study |  |
| YJH682 | WT | *MATa leu2-3 112 trp1-1 can1-100 ura3-1 ade2-1 his3-11,15* | Jay Hesselberth | 10,11 |
| YJH835 | *trl1* | *MATa leu2-3 112 trp1-1 can1-100 ura3-1 ade2-1 his3-11,15 trl1::KanMX (pAG424-10x-tRNA)* | Jay Hesselberth | 10,11 |
| YLB313 | *dom34* | *MATa leu2-3 112 trp1-1 can1-100 ura3-1 ade2-1 his3-11,15 dom34::NatMX* | this study |  |
| YLB317 | *trl1 dom34* | *MATa leu2-3 112 trp1-1 can1-100 ura3-1 ade2-1 his3-11,15 trl1::KanMX (pAG424-10x-tRNA) dom34::NatMX* | this study |  |

**Supplementary Table 2.**
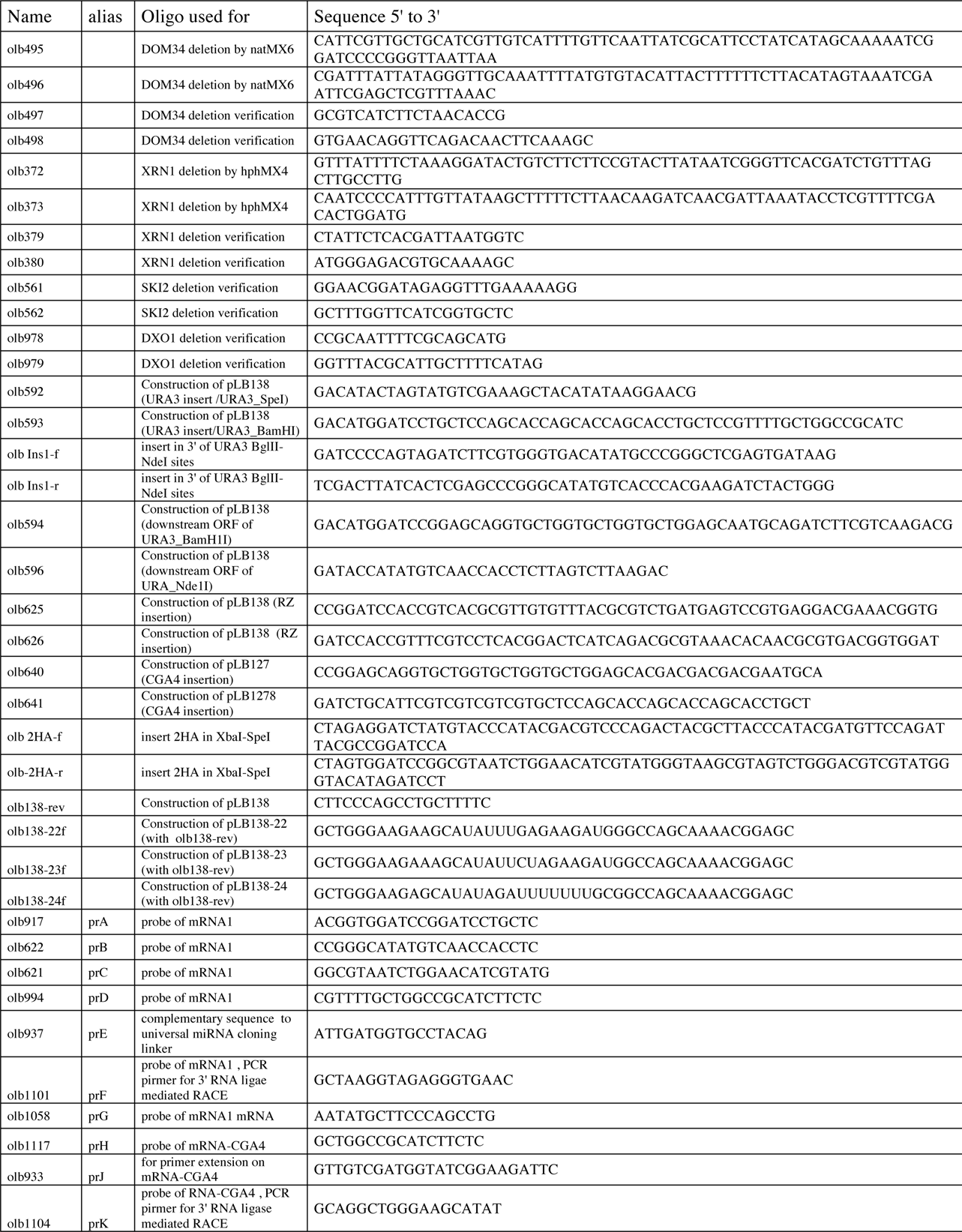
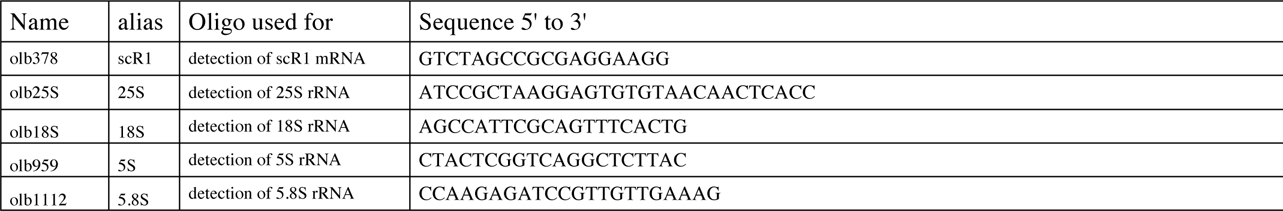
Oligonucleotides used in this study

**Supplementary Table 3.**
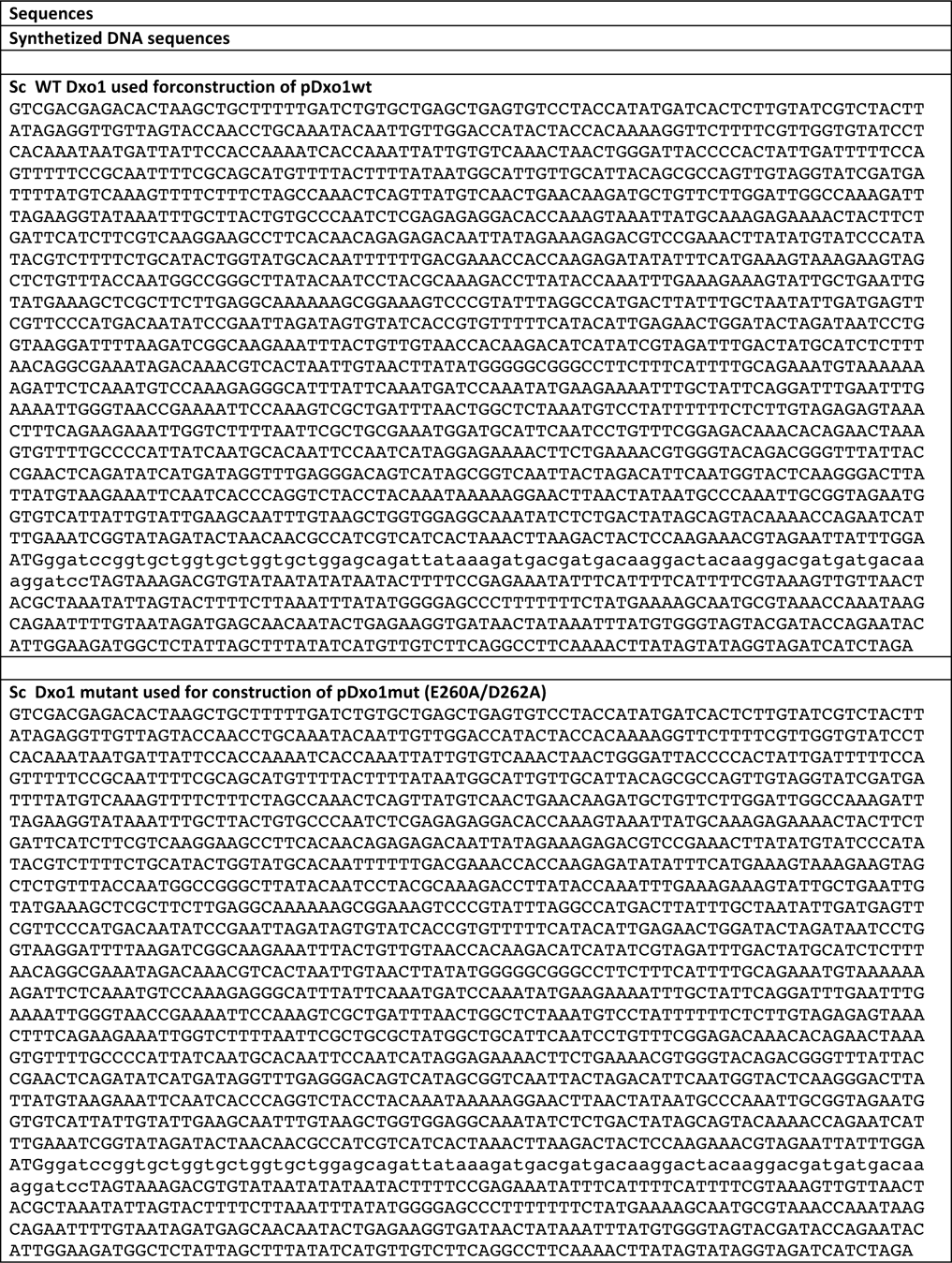
Synthetized DNA DXO1 sequences for plasmid constructions

**Supplementary Table 4.** Plasmids used in this study

| Plasmid | Description | Marker | Reference |
| --- | --- | --- | --- |
| p138 | expression of RNA1Rz | LEU2 | This study |
| p127 | expression of (CGA)4-mRNA | LEU2 | This study |
| p127T | expression of (CGA)2-mRNA | LEU2 | This study |
| pDxo1WT | expression of Dxo1 of *S. cerevisiae* from pRS313 plasmid derivative | HIS3 | This study |
| pDxo1mut | expression of a catalytic mutant of Dxo1 of *S. cerevisiae* from pRS313 plasmid derivative | HIS3 | This study |
| pFA6a-natMX6 | natMX6 confers resistance to nourseothricin |  | 5 |
| pAG32 | hphMX4 confers resistance to hygromycin B |  | 6 |
| pRS316 | Centromeric | URA3 | 8 |
| pRS313 | Centromeric | HIS3 | 8 |

## Notes

#### Summary of Updates

Updated version with detailed information in the Methods section and clarified Figures 1f and 5b

